# Integrative Nanopore and Illumina sequencing reveals age-associated tRNA modification and CCA-tail dynamics in yeast

**DOI:** 10.64898/2026.09.22.753482

**Authors:** Katherine M Brazell, Christopher Prevost, Natalia Zawrotna, Carrie Smith, Lindsey N Power, Jeffrey S Smith, Zhangli Su

## Abstract

Aging is characterized by a progressive loss of proteostasis. Transfer RNAs (tRNAs) are essential regulators of translation, yet their dynamics during aging remain poorly understood due to challenges in sequencing highly modified RNAs. Here we present a benchmarked Nanopore direct RNA sequencing (RNA004 chemistry) resource that profiles the *Saccharomyces cerevisiae* tRNAome during replicative aging at single-molecule resolution. Using *in vitro* transcribed tRNA controls, we establish modification detection thresholds and validate key findings with orthogonal Illumina sequencing. While overall tRNA abundance remains largely stable, our resource reveals age-associated terminal A cleavage at the 3’ CCA tail of mature tRNAs, targeted T-loop and anticodon modification changes, and single-molecule evidence of modification co-occurrence. This dataset provides a resource for exploring tRNA regulation, translation fidelity, and longevity.

## INTRODUCTION

Aging is a complex biological process of progressive physiological decline. This decline features distinct hallmarks, with the loss of proteostasis being a critical driver of age-related pathologies.^1^ Proteostasis is maintained by the equilibrium between protein synthesis, folding, function and degradation. As organisms age, this intricate balance is compromised – the capacity to accurately synthesize and fold proteins diminishes, resulting in the accumulation of misfolded proteins and potentially toxic aggregates.^2–4^ Other hallmarks of aging can compound loss of proteostasis, worsening the proteotoxic burden and accelerating aging in a vicious cycle. In a seemingly short-term strategy to mitigate the proteotoxic stress, cells respond by down-regulating global protein synthesis.^5–7^ However, specific proteins in pro-adaptation pathways are also up-regulated through translation.

Translational regulation relies on transfer RNAs (tRNAs), which are essential adaptor molecules for protein synthesis and cellular homeostasis. In recent years, there has been a shift in our understanding about tRNA molecules being dynamic regulators of translation programs in diverse biological contexts. tRNAs are historically difficult to profile due to their highly modified nature and strong secondary structure. Recent advancements in high-throughput sequencing techniques, especially Nanopore tRNA-sequencing, have greatly facilitated this profiling.^8,9^ Traditional Illumina sequencing methods often require reverse transcription into cDNA, which can be inefficient for highly structured and modified RNAs, such as tRNA. Nanopore allows for the direct sequencing of the RNA molecule and preserves full-length tRNA sequences, including their post-transcriptional modifications.^8–10^ This capability enables the accurate quantification of individual tRNAs and the detection of their modification patterns.

Previous literature supports a possible link between tRNA regulation and aging. For example, inhibiting the RNA polymerase III to reduce tRNA synthesis has been shown to extend lifespan in yeast, worms and flies.^11,12^ Similarly, the deletion of *LOS1*, the nuclear exporter of tRNA, also prolongs lifespan in both yeast and worms.^13^ Recent studies have begun to address this gap: METTL1-mediated m^7^G tRNA modifications were shown to decline during mammalian senescence, and age-dependent loss of manQ modification has been linked to translational fidelity changes in 2BS cells and mouse plasma.^14,15^ However, these studies focused on individual modification types, and no comprehensive, global atlas of tRNAome-wide expression and modification dynamics during aging currently exists.

Here we present a quantitative resource of the *Saccharomyces cerevisiae* tRNAome across replicative aging using Nanopore direct RNA sequencing at single-molecule resolution, complemented by Illumina-based Induro-tRNA-seq. We benchmarked our pipeline and established an *in vitro* transcribed (IVT) tRNA control to define accurate error thresholds for endogenous modification detection, optimizing classification through error component decomposition and neighbor-effect smoothing. Systematic comparison of Nanopore and Illumina platforms reveals complementary modification detection profiles, with Nanopore capturing modifications invisible to reverse transition-based methods. While overall tRNA abundance remains largely stable during aging, our atlas reveals systematic remodeling of the tRNAome. A global loss of the terminal 3’ CCA nucleotide of mature tRNAs is consistent with enzyme-mediated cleavage upon aging stress. T-loop modification signatures increase in specific tRNA species including tRNA^Lys-CTT^, and anticodon loop m^3^C_32_ modification shows age-dependent elevation. Finally, we found that co-occurrence of modifications on individual tRNA molecules is disrupted in aged cells. Together, this resource provides a comprehensive framework for studying tRNAome dynamics during aging.

## RESULTS

### Global profiling of tRNA expression and modifications using Nano-tRNA-seq

We utilized the Nano-tRNA-seq method^8^ to identify global tRNA changes, which captures mature tRNAs with their CCA-end via a splint adaptor (Figure 1A). This approach allows for the sequencing of native RNA molecules and can measure tRNA abundances and chemical modifications to the tRNA without reverse transcription and amplification. We first compared our dataset on biological tRNAs isolated from *Saccharomyces cerevisiae* with previously published tRNA-seq studies. Quantification of tRNA abundance revealed strong concordance between our results and those previous studies using Nanopore^8^ and Illumina^16^ platforms (Figure S1A-B). This indicates that our sequencing pipeline reliably captures tRNA expression levels. It has been shown that tRNA modifications can be captured as Nanopore base-calling error signatures (including mismatch, insertion, and deletion) due to how they chemically differ from the canonical bases.^8,9,17,18^ As a proof of concept, we quantified relative m^1^A (*N*1-methyladenosine) level based on error percentages at the known m^1^A sites^19^ in our study and two independent datasets. The error percentage at m^1^A sites in our dataset was significantly correlated with both *Lucas et al.* 2023^8^ and *Behrens et al.* 2021^16^ (Figure S1C-D), demonstrating strong feasibility of using base-calling errors to quantify tRNA modifications by Nanopore. Crucially, because each Nanopore read represents a single tRNA molecule, our resource enables single-molecule analysis of modification patterns, including per-read stoichiometry and combinatorial modification states across individual tRNA molecules.

**Figure 1.**
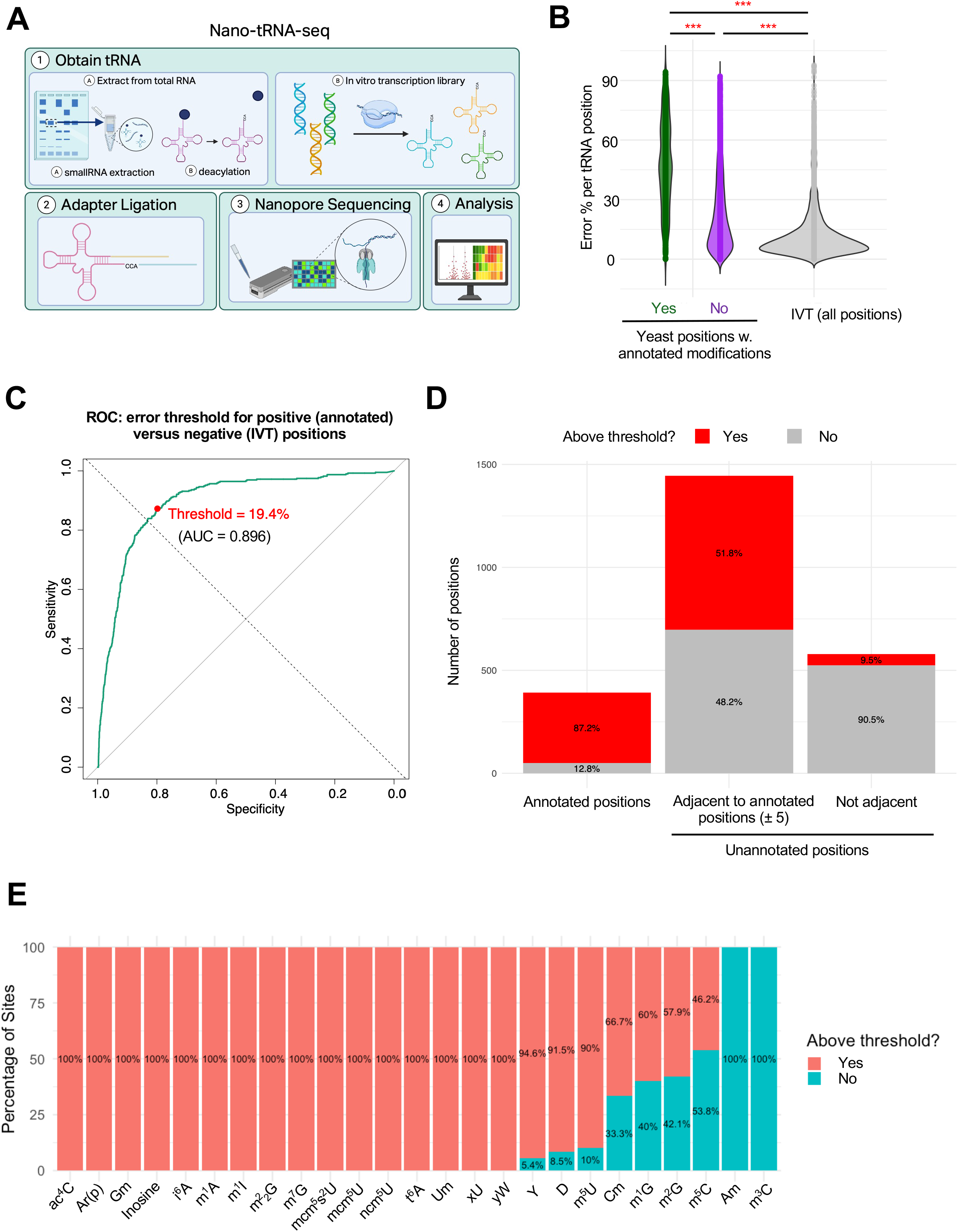
Global profiling of tRNA expression and modifications using Nano-tRNA-seq. (A) Schematic overview of the experimental method for Nanopore tRNA sequencing across biological and *in vitro* transcribed (IVT) tRNAs. (B) Comparison of global total error rates between yeast and IVT tRNAs. Yeast error rates are separated into sites corresponding to canonically annotated modifications (based on MODOMICS) and sites without annotated modifications. Statistical significance was evaluated using Student’s t-test. (C) Receiver Operating Characteristic (ROC) curve plots the true positive rate against the false positive rate for varying score thresholds. Annotated yeast modification site error rates were designated as true positives, while IVT sites were designated as true negatives. The area under the curve (AUC) is 0.896 with a sensitivity of 0.872449 and specificity of 0.7983574. Youden’s threshold was calculated to be 0.194. (D) Number of modification sites with error rates above or below the IVT baseline. Sites with error rates exceeding the IVT rate are highlighted in red. (E) Percentage of occurrences in which each specific modification site exhibits an error rate at or above the IVT baseline.

*Saccharomyces cerevisiae* transcribes 54 unique mature nuclear-encoded tRNA sequences in total, with 12 sequences being highly similar (having 95-99% identity). To ensure reliable quantification, we kept the 42 remaining sequences, out of which only 31 tRNAs have been annotated by MODOMICS for base modifications.^8,19^ To identify the optimal threshold for calling tRNA base modifications, especially on the previously unannotated tRNAs, we needed a modification-free control to distinguish sequencing errors arising from true RNA modifications versus those due to background noise or technical artifacts.^8,20^ To this end, we *in vitro* transcribed (IVT) all 54 unique mature tRNA sequences lacking endogenous modifications (Figure 1A). The IVT tRNA library is designed to end with CCA-end following BsaI-HFv2 digestion to mimic the endogenous tRNAs. The integrity and purity of IVT tRNAs were confirmed by urea-PAGE analysis, which showed the expected sizes and digestion patterns (Figure S1E). As expected, IVT tRNAs exhibited a low baseline error percentage, with only a small fraction of positions exhibiting elevated background errors (Figure 1B). Global error analysis revealed that yeast tRNAs displayed significantly higher error percentage per position compared to IVT tRNAs at canonically annotated modification sites (Figure 1B). In contrast, sites without annotated modifications in yeast tRNAs (median 17.2%) showed lower error percentage than annotated sites (median 47.4%), but still higher than the IVT baseline (median 8.82%). Receiver-Operator Curve (ROC) analysis of true positive (sites annotated by MODOMICS) and true negative sites (IVT sites) identified error threshold of 19.4% to maximize area under the curve, with sensitivity of 87.2% and specificity of 79.8% (Figure 1C).

Using this error threshold, we next quantified the number of modification sites with error rates either above or below the threshold. A substantial subset (87.2%) of annotated modification sites displayed error rates more than 19.4% (Figure 1D, left), higher than unannotated positions. These high-error sites are highlighted in red. Finally, we assessed whether the percentage of sites passed the error threshold is affected by the chemical nature of the modification type (Figure 1E). This analysis revealed that 100% of known modification types displayed elevated error rates. These include m^1^A, m^7^G (*N*7-methylguanosine), ac^4^c (*N*4-acetylcytidine), Ar(p) (2’*O*-ribosyladenosine), Gm (2’*O*-methylguanosine), i^6^A (*N*6- isopentenyladenosine), m^1^I (1-methylinosine), m^2^_2_G (*N*2,*N*2-dimethylguanosine), mcm^5^s^2^U (methoxycarbonylmethyl-2-thiouridine), mcm^5^U (5-methoxycarbonylmethyluridine), ncm^5^U (5- carbamoylmethyluridine), t^6^A (*N*6-threonylcarbamoyladenosine), Um (2’-O-methyluridine), xU (unknown modified uridine), yW (wybutosine), and inosine. This is consistent with the presence of endogenous modifications that alter ionic current signals and base-calling accuracy during Nanopore sequencing. On the other hand, we also noticed that tRNA modification types like m^3^C (*N*3-methylcytidine), Am (2’*O*- methyladenosine), m^5^C (5-methylcytidine), m^2^G (*N*2-methylguanosine), m^1^G (1-methylguanosine), Cm (2’*O*-methylcytidine) have a significant portion of sites that fall below the threshold. This suggests either the Nanopore error threshold could not robustly capture these modification types, or the endogenous modification stoichiometry is low. Interestingly, focusing on the 31 tRNAs with annotated modifications, we also identified unannotated sites within yeast tRNAs that exhibited error rates exceeding the error threshold (Figure 1D, right). Consistent with the idea that the pore reads a stretch of sequence at a time, positions adjacent to annotated modification positions (defined as ± 5 nucleotides) have higher error than positions not adjacent to modification sites (Figure 1D, right). This highlights the importance of considering the effect of modification at neighboring positions.

Because Nanopore basecalling errors comprise mismatch, insertion and deletion, we also tested whether each component individually separates modified from unmodified positions. ROC analysis revealed that mismatch alone is nearly as informative as total error (AUC = 0.91 vs 0.93), while insertion and deletion are weaker discriminators (Figure S1F). We further tested whether accounting for the known neighbor effect of Nanopore modifications improves classification. Applying a ± 1, ± 2 and ± 3-nucleotide sliding window to the error rate substantially improved ROC performance, raising the AUC from 0.93 to 0.96 (Figure S1G). Altogether, these benchmarking analyses confirm that Nanopore direct RNA sequencing discriminates known tRNA modifications from unmodified positions with high sensitivity and specificity, validating the approach for profiling modification dynamics across age conditions.

### Nano-tRNA-seq reveals modification-specific error signatures complementary to Illumina-based detection

We systematically characterized basecalling error profiles at annotated tRNA modification sites using Nano-tRNA-seq. Decomposition of Nanopore errors into mismatches, insertions, and deletions at ± 2 positions revealed modification-specific error type compositions (Figure 2A). m^1^A and m^2^_2_G showed high proportions of both mismatches (35% and 32%) and deletions (37% and 36%), while m^5^U was dominated by deletions (27%) with relatively low mismatches (10%). Pseudouridine (Ψ) showed primarily mismatches (37%) with moderate deletions (17%), and D modifications showed balanced contributions across all three error types. Five additional modification types (t^6^A, ac^4^C, I, Cm, i^6^A) exhibited similar patterns (Figure S2A).

**Figure 2.**
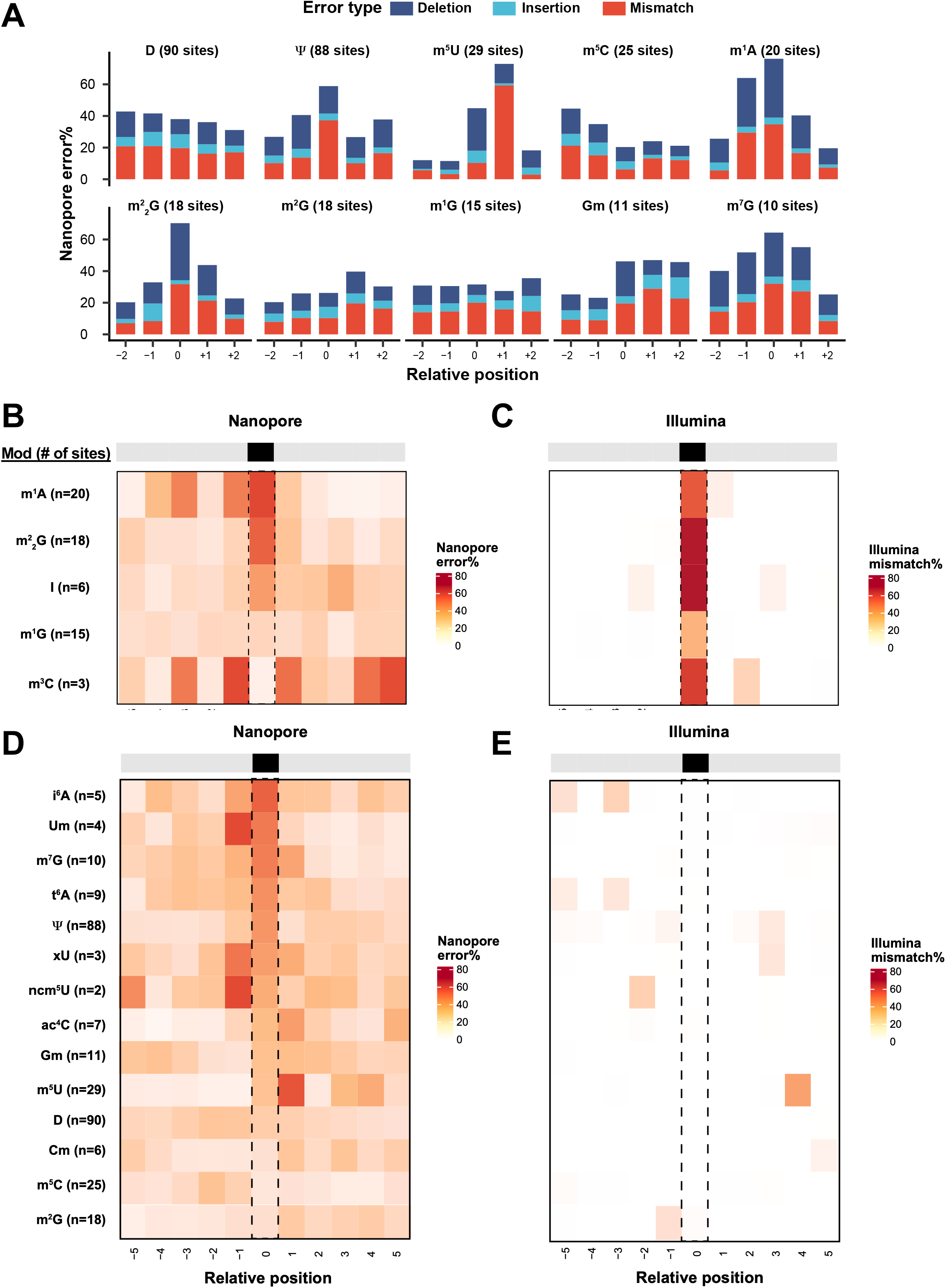
Nano-tRNA-seq reveals modification-specific error signatures in yeast tRNAs. (A) Error type composition (mismatch, insertion, deletion) for the 10 most prevalent modification types by number of annotated sites. Bars show the mean proportion of each error type within ±2 positions of modification sites, averaged across all instances per modification type. (B-E) Heatmaps showing position-specific mismatch rates across ±5 positions surrounding annotated modification sites, for Nanopore (B and D) and Illumina (C and E). Each row represents a modification type; columns represent sequential positions relative to the modification site (0). Color scale spans 0 to the maximum observed mismatch rate per platform.

Next, we compared Nanopore error profile to Illumina-based profiles. For the Illumina arm, we employed Induro reverse transcriptase (RT), a group II intron-encoded RT with high processivity that enables efficient read-through of tRNA modifications. Focusing on the seven known modification types with detectable Illumina RT signatures (m^1^A, m^2^_2_G, Inosine, m^1^G, m^3^C, m^1^I and yW), we computed position-specific error rates across ± 5 positions surrounding each modification site (Figure 2B-C and Figure S2B-C). Illumina produced sharp base resolution with low background (∼0.1%) and successfully detected strong mismatch signals at the modification site (position 0) for all modification types. For this group of modifications, Nanopore error peaks at the m^1^A, m^2^_2_G, I (Inosine), m^1^I and yW modification sites, but not at m^1^G and m^3^C sites. m^3^C is particularly interesting, as its Nanopore error peaks at −1 position, which is not annotated with any modifications. We attribute this to the basecaller correctly identifying m^3^C as C while the modification perturbs the k-mer signal sufficiently to induce errors at the flanking nucleotide.

For the modification types that do not disrupt Watson-Crick base pairing and are not detectable by Illumina RT signatures (Figure 2D-E and Figure S2B-C), Nanopore detected a broader range of modifications at the modification site (position 0), including i^6^A, t^6^A, m^7^G and pseudouridine (Ψ), all of which were invisible to Illumina (<1% mismatch rate). Additional modification types exhibited substantial Nanopore signals but with peaks at neighboring positions rather than at the modification sites, likely reflecting the multi-nucleotide k-mer model of the basecaller and/or contamination from adjacent modifications. For example, m^2^G showed a prominent peak at position −1 (26% mismatch) rather than at position 0 (3%). This signal arises from m^1^G at the neighboring position. The m^1^G_9_/m^2^G_10_ pair present in 6 of 18 m^2^G sites produced an RT signature that extends to the adjacent nucleotide. This complementarity highlights the advantage of Nanopore direct RNA sequencing for modifications that do not perturb reverse transcription. Consistent with recent report,^21^ we noted some modifications including D, m^5^U and m^5^C do not show a peak at the modification sites by Nanopore. The +1 signal from m^5^U is likely from Ψ55, the universally conserved T-loop pseudouridine present at +1 in 28 out of 29 m^5^U sites.

Overall, Nanopore error profiles displayed elevated background across the entire ± 5 window (20-40% mismatch at flanking positions), reflecting the multi-nucleotide k-mer model used by the basecaller, whereby modifications influence the signal of neighboring k-mers.^18^ Illumina profiles, by contrast, showed minimal background with signals largely confined to the modification site. Neighboring modifications can contaminate error profiles at adjacent positions, for example, Ψ showed elevated signal at position −1 (8%) in Illumina, attributable to neighboring modifications in the T-loop. The m^2^G −1 signal from neighboring m^1^G, discussed above, provides another example. Together, these results demonstrate that Nanopore and Illumina exhibit complementary detection profiles, with Nanopore capturing modifications invisible to RT-based methods and Illumina providing sharper positional resolution for modifications that disrupt Watson-Crick base pairing.

### tRNA expression is stable during aging while CCA tails undergo cleavage

Replicative aging in *Saccharomyces cerevisiae* provides a well-established model for studying age-associated molecular changes and is defined by the number of daughter cells a mother cell produces before senescence.^22,23^ In our study, young cells were defined as the logarithmically growing cultures prior to aging isolation. We aged the experimental cells for 36 hours. This specific timeframe allows mother cells to reach a sufficiently advanced replicative age where proteostasis stress is pronounced, consistent with established methodologies.^24^ To examine how tRNA abundance changes with replicative aging, we quantified tRNA expression across young and old *Saccharomyces cerevisiae* replicative aging samples by both Nanopore and Illumina sequencing. Overall, both platforms show high concordance in capturing tRNA expression, with a Pearson correlation R of 0.77 (Figure 3A). DESeq2-normalized tRNA levels were largely consistent between age groups, with only tRNA^Tyr-GTA^ exhibiting a mild but significant decrease in expression in older samples by both platforms (Figure 3B). Each platform also captures unique tRNAs that show mild by significant changes (Data S1).

**Figure 3.**
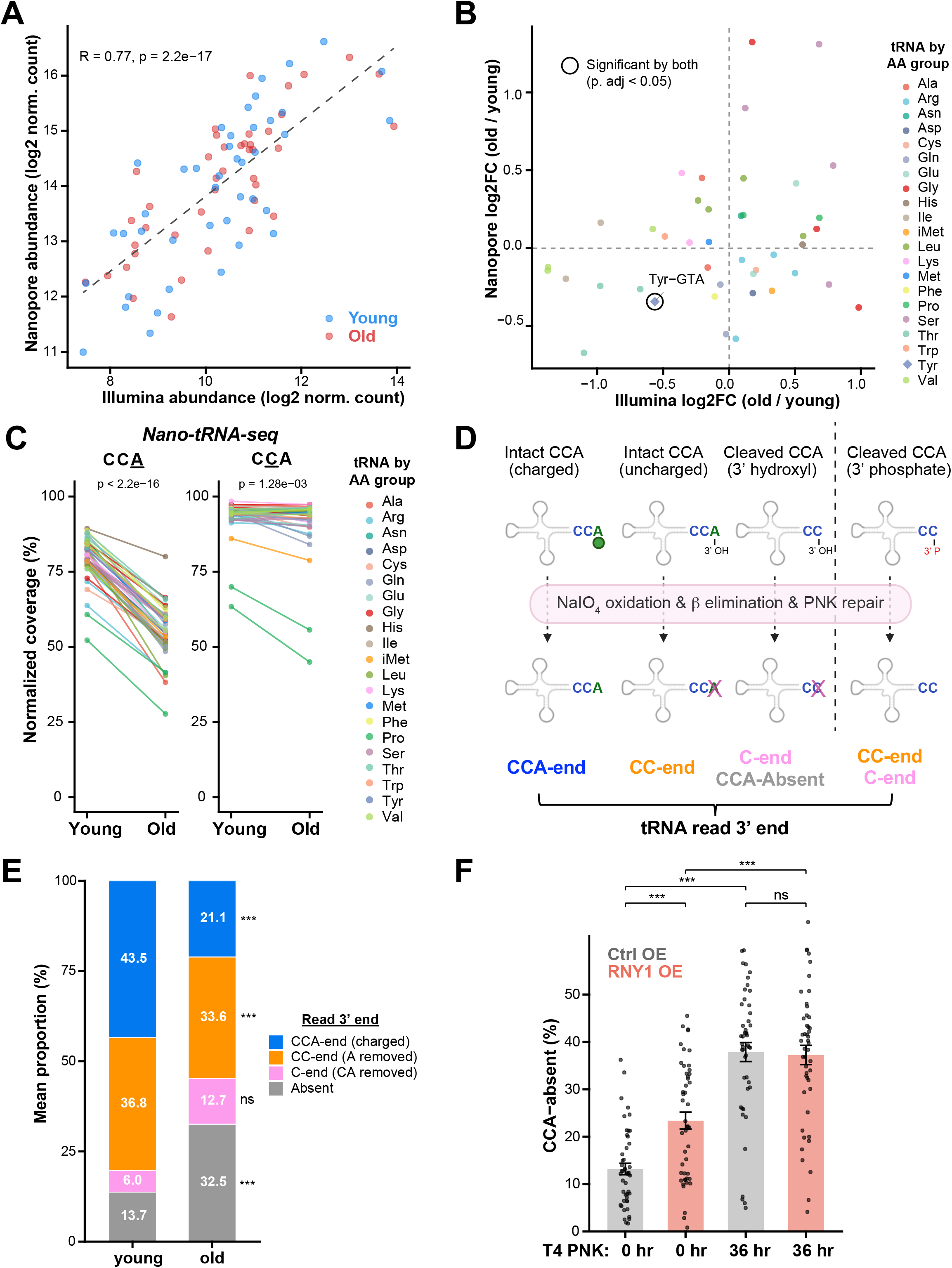
tRNA expression is stable during aging while CCA tails undergo cleavage. (A) Correlation of tRNA expression between Nanopore and Illumina platforms (Pearson correlation; each point represents one cytoplasmic tRNA at anticodon level). (B) Log2 fold change (old/young) correlation between platforms. tRNAs are colored by amino acid family, differentially expressed tRNAs (DESeq2, p. adj < 0.05) on both platforms labeled. (C) Nano-tRNA-seq detects 3’ CCA integrity loss. Normalized coverage from the terminal A or second C to the first C in the CCA tail across different tRNAs. Each line represents one cytoplasmic tRNA at anticodon level. Statistics by paired t-test. (D) Schematic of the charging protocol to dissect 3’ end dynamics with Illumina pipeline. (E) 3’ end composition in young versus old samples by charging protocol. Stacked bars show mean proportions across cytoplasmic tRNAs. (F) CCA-absent fraction in Rny1 OE versus control by T4 PNK Illumina analysis. Bars show mean proportions across cytoplasmic tRNAs with individual values overlaid. (E-F) Statistics by paired t-test with Bonferroni correction (*** p < 0.001, NS p > 0.05).

During the analysis, we noticed an aging-dependent drop in coverage at the terminal A position of the CCA tail across several tRNAs by Integrative Genomics Viewer (IGV) visualization (Figure S3A). The CCA tail is essential for tRNA aminoacylation and proper translation, and cleavage or damage to this sequence could cause changes in tRNA stability or function.^25^ We systematically quantified the tRNA 3’-end integrity by measuring the coverage at each position of the CCA tail. Quantification of normalized coverage from the terminal A to the first C confirmed that, while full-length tRNAs were generally well represented, older samples displayed a pronounced reduction in coverage at the terminal A position (Figure 3C). This effect appeared specific to the terminal A, as coverage of the second C of the CCA tail showed no appreciable difference between young and old samples (Figure 3C). The Nanopore protocol uses a splint adaptor to recognize the 3’ CCA end on mature tRNAs, which limits resolution of the 3’ end dynamics. We therefore adopted the standard charging protocol for our Illumina pipeline, which includes periodate oxidation and beta-elimination, followed by PNK repair (Figure 3D). Specifically, charged tRNAs are protected from oxidation by the aminoacyl group and retain full CCA ends, whereas uncharged tRNAs with free 3’ hydroxyl group undergo beta-elimination of the 3’ terminal nucleotide. Consequently, reads retaining a CCA end indicate charged tRNAs, while CC-end, C-end or CCA-absent reads represent uncharged tRNAs with varying degrees of 3’ end degradation. Using this protocol, we confirmed age-dependent 3’ CCA integrity loss by Illumina sequencing (Figure 3E): the charged fraction (CCA-end reads) is significantly reduced in old samples, while CCA cleavage is significantly increased, as indicated by the elevated fraction of C-end and CCA-absent reads. The full 3’ end repertoire per tRNA can be explored in Data S2. The Illumina confirmation suggests Nanopore is capable of detecting tRNA 3’ end loss.

tRNA 3’ end CCA cleavage has been reported in the case of oxidative stress, but the mechanism and responsible enzyme have not been well understood.^8,26^ Since Rny1 has been previously described to cleave mature tRNAs at the anticodon loop upon oxidative stress, we tested whether Rny1 over-expression can cleave the 3’ CCA end of mature tRNAs. Rny1 belongs to the RNase T2 family, which produces 3’ phosphate termini via a 2’,3’ cyclic phosphate intermediate; these termini are resistant to periodate oxidation and therefore cannot be directly detected by the charging protocol. We therefore performed T4 PNK treatment to convert 3’ phosphate and cyclic phosphate to 3’ hydroxyl group before Illumina library preparation. We found Rny1 over-expression significantly increased both C-end and CCA-absent fractions in young cells (Figure 3F, Figure S3B), with a corresponding reduction in CCA-intact reads (Figure S3C), demonstrating that Rny1 is capable of cleaving the tRNA 3’ CCA end. Interestingly, Rny1 over-expression in aged cells increased the C-end fraction but did not further elevate the CCA-absent fraction (Figure 3F, Figure S3B-D), suggesting that aging has already depleted the pool of tRNAs susceptible to complete CCA removal and the remaining CCA-containing tRNAs are only partially accessible to Rny1, yielding C-end rather than CCA-absent products. Altogether, these results identify Rny1 as a candidate nuclease sufficient to cleave tRNA 3’ CCA ends and support a model in which age-dependent Rny1 activation contributes to the progressive CCA tail loss during replicative aging.

### Replicative aging alters specific tRNA T-loop modification signatures

Base modifications on tRNAs are key regulators to tRNA stability and translation fidelity.^27–29^ As demonstrated above, Nanopore direct RNA sequencing detects diverse tRNA modification types through position-specific error signatures (Figure 1). To investigate how tRNA modification patterns change with age, we compared Nanopore sequencing error rates between young and old *Saccharomyces cerevisiae* replicative aging samples. Analysis of directional changes in error rates across all tRNA positions revealed a subset with significantly altered error frequencies between age groups (Figure S4A). Using canonical tRNA coordinates for consistent numbering, metagene analysis across 42 cytoplasmic tRNAs showed that aging increased error preferentially around the T-loop region (corresponding to positions 54 to 60), with positions 57 and 58 showing the strongest enrichment for both error magnitude and number of significant tRNAs (Figure 4A). Because Nanopore basecalling integrates signal from multiple consecutive bases, error signals at modification sites can spread to neighboring positions; the smoothed line (± 3 positions) accounts for this signal spread. Visualization on a tRNA canonical cloverleaf structure confirmed that T-loop positions display the largest age-dependent error increases without smoothing (Figure 4B). Notably, position 57 has no annotated modification but likely reflects error spillover from adjacent m^1^A_58_ (Figure S4B).

**Figure 4.**
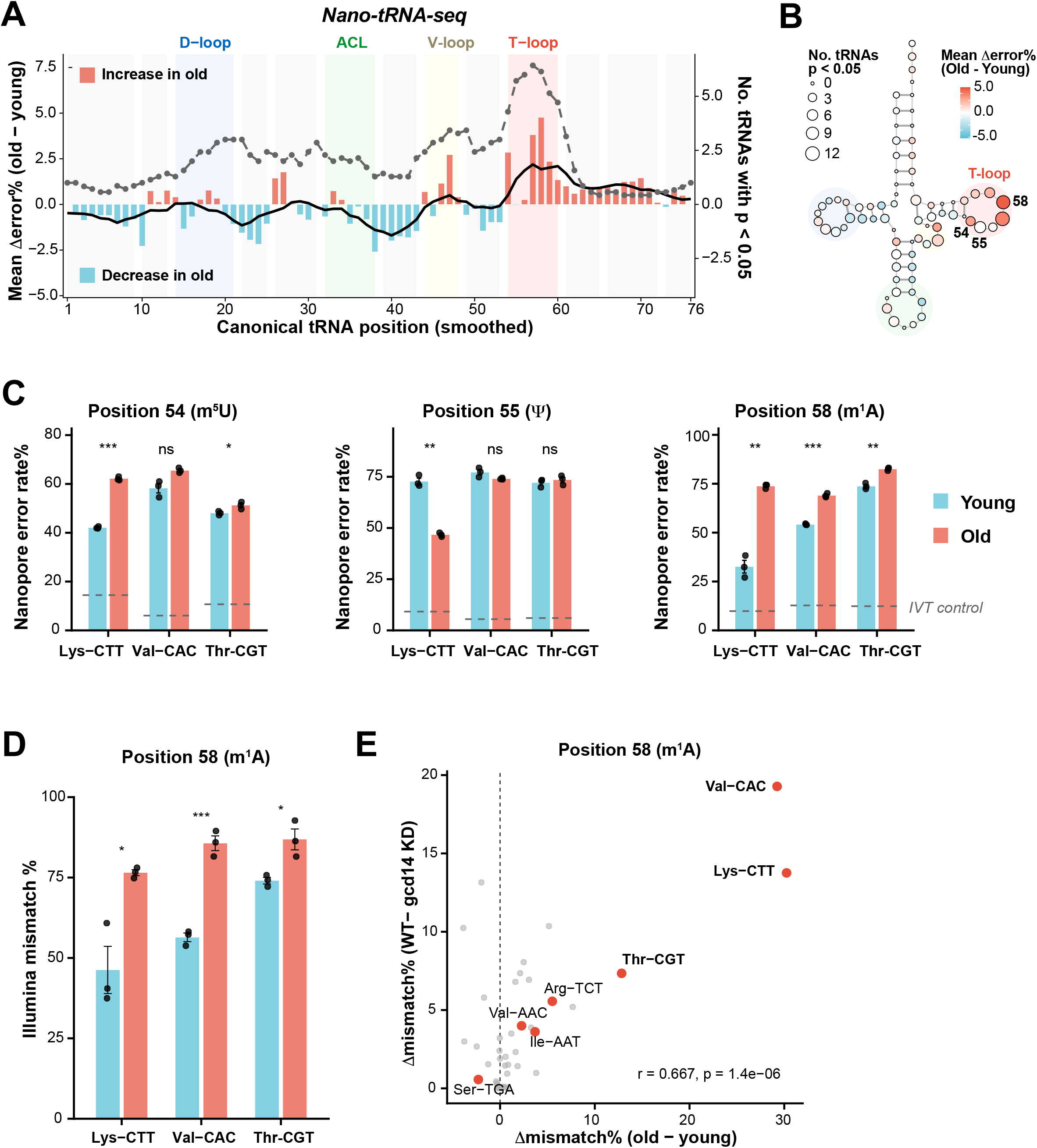
Replicative aging alters specific tRNA T-loop modification signatures. (A) Metagene analysis of age-dependent Nanopore error changes across tRNA positions highlights T-loop. Bars show mean error change (old – young) per position across 42 tRNAs; red indicates increased error in old, blue in young. The black line shows the smoothed (±3 positions) mean error change. The dashed grey line (right axis) shows the smoothed (±3 positions) number of tRNAs with a significant change (two-sided t-test, p < 0.05). Structural regions are shaded: D-loop (blue), ACL or anticodon loop (green), variable loop (yellow), T-loop (pink). (B) tRNA cloverleaf heatmap. Each circle represents one tRNA canonical position, colored by mean error change (old - young) and sized by the number of tRNAs with a significant change (p < 0.05). Structural regions are shaded as in (A). (C) Bar plots show mean Nanopore error rate (± SEM, n = 3 biological replicates) for tRNA^Lys-CTT^, tRNA^Val-CAC^, and tRNA^Thr-CGT^ at positions 54 (m^5^U), 55 (Ψ), and 58 (m^1^A). Blue bars = young, red bars = old; black dots = individual replicate values. Dashed grey lines indicate the IVT control error rate (unmodified tRNA baseline) for each tRNA. Significance by two-sided Student’s t-test: * p < 0.05, ** p < 0.01, *** p < 0.001. (D) Bar plots show mean Illumina mismatch rate (± SEM, n = 3 biological replicates) for tRNA^Lys-CTT^, tRNA^Val-CAC^, and tRNA^Thr-CGT^ similar to panel (C). (E) Correlation between age-dependent and *GCD14*-dependent m^1^A_58_ changes. Each point represents one of 42 cytoplasmic tRNAs; x-axis shows aging-dependent mismatch% change (old – young), y-axis shows *GCD14*-dependent mismatch% change (WT – *GCD14* KD). Significant tRNAs are colored in red and labeled (two-sided t test, p < 0.05).

To further understand the specificity of T-loop changes, we quantified error rates at T-loop modification positions 54 (m^5^U), 55 (Ψ) and 58 (m^1^A) for the three tRNAs showing the strongest age-dependent changes in T-loop, tRNA^Lys-CTT^, tRNA^Val-CAC^ and tRNA^Thr-CGT^ (Figure 4C). tRNA^Lys-CTT^ in aged samples displayed the most significant change across all three positions, with m^5^U and m^1^A error significantly exceeding young samples, while Ψ error was lower in old samples (Figure 4C, Figure S4B). Notably, this change is specific to tRNA^Lys-CTT^, but not to tRNA^Lys-TTT^, whose sequence differs by 20 nt (Figure S4C-D), suggesting specific regulation. In addition to tRNA^Lys-CTT^, tRNA^Val-CAC^ and tRNA^Thr-CGT^ also showed significant m^1^A_58_ increase by Nanopore (Figure 4C). The m^1^A increase is further validated by Illumina-based mismatch analysis, which confirmed significant increases in the same three tRNAs (Figure 4D and Figure S4E).

m^1^A_58_ is catalyzed by the yeast Gcd10/Gcd14 complex, with Gcd14 as the catalytic subunit. We reasoned that the significant increase in m^1^A on multiple tRNAs could be due to altered activity of the complex. Supporting this hypothesis, we sequenced *GCD14* knockdown samples and found a strong correlation between age-dependent m^1^A increase and *gcd14* KD-dependent m^1^A loss (Figure 4F). This suggests that the tRNAs most affected by aging are also most sensitive to *GCD14* dosage, supporting a model where age-dependent m^1^A changes reflect altered Gcd14 complex activity.

### Age-dependent m^3^C_32_ increase and altered modification co-occurrence

While Nanopore basecalling errors robustly detect T-loop modifications (Figure 4), certain modification types produce mismatch signatures that are better resolved at single-base resolution by Illumina. Illumina detects modifications at positions 9, 26, 32, 34, 37 and 58 with high mismatch rates (Figure S5A). Because Figure 1E established that Nanopore basecalling cannot capture all tRNA modifications, we examined the Illumina mismatch landscape across all canonical positions for additional age-dependent changes beyond the T-loop. This revealed position 32 in the anticodon loop as the second-strongest age-dependent signal after position 58 (m^1^A_58_), with significant mismatch increases in six tRNAs (Figure 5A and Figure S5B). Since Illumina has tight error distribution (Figure 2), no smoothing was applied. Position 32 carries m^3^C, installed by TRM140 on Ser and Thr tRNAs. Among the four known m^3^C_32_ substrates (Ser-TGA, Ser-GCT, Thr-TGT, Thr-AGT), all show significant age-dependent mismatch increases, with Ser tRNAs exhibiting larger effects (by 14-24%) than Thr tRNA (by 2.7-3.6%) (Figure 5B). Consistent with Figure 2B, Nanopore basecalling errors at position 32 remain near background for all four tRNAs (0.01-0.61%) but shows significant increase at position 31 for tRNA^Ser-TGA^ (Figure 5C). Together, these results demonstrate that Nanopore can detect aging-associated m^3^C changes as error shifts at the adjacent upstream position but with lower sensitivity than Illumina.

**Figure 5.**
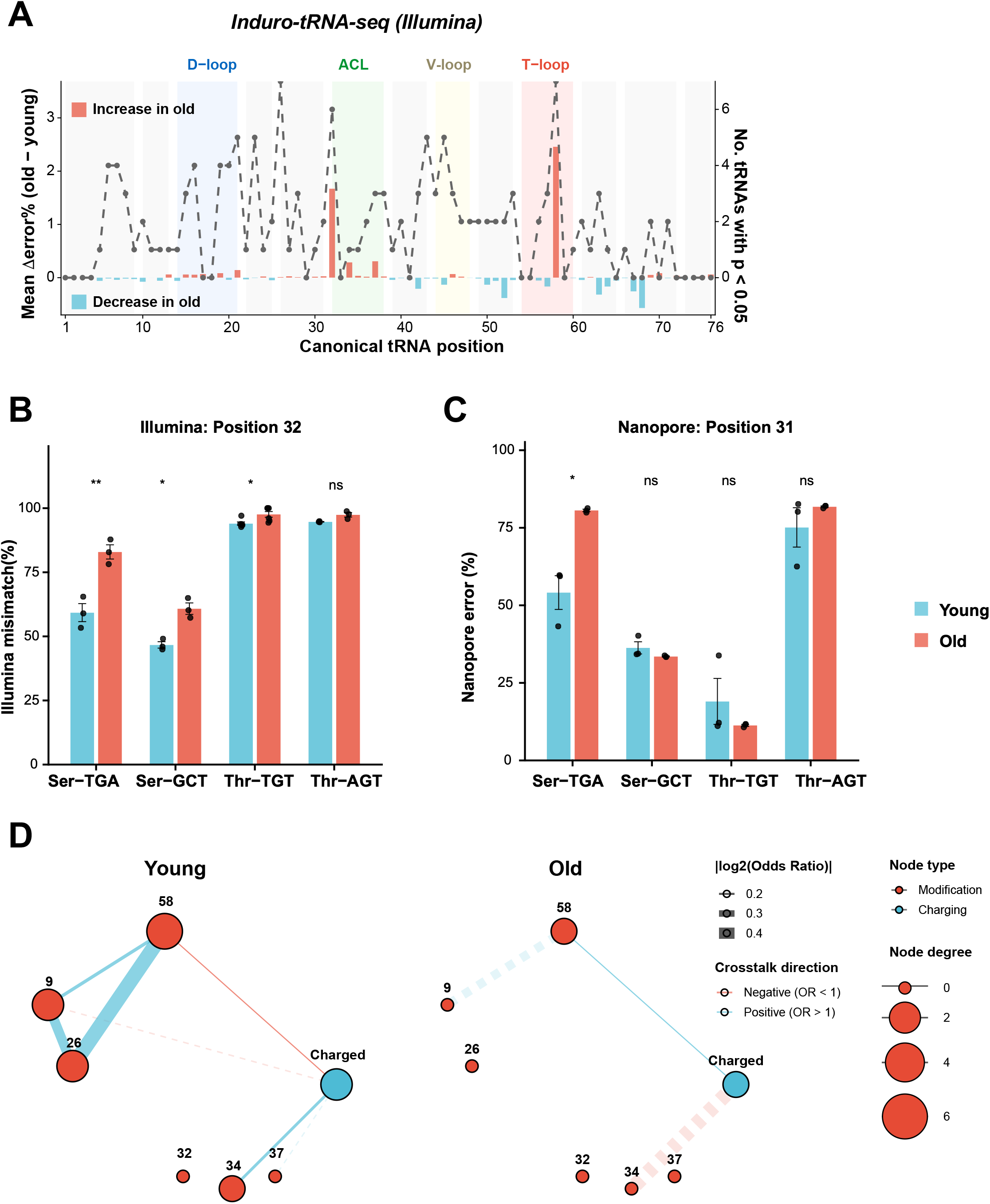
Age-dependent m^3^C_32_ increase and altered modification co-occurrence. (A) Metagene profile showing per-position mismatch rates from Illumina Induro-tRNA-seq across all 76 canonical tRNA positions for young (blue) and old (red) yeast samples, similar to Figure 3A but without smoothing. (B-C) m^3^C modification levels at position 32 for four known m^3^C substrate tRNAs (Ser-TGA, Ser-GCT, Thr- CGT, Thr-AGT) measured by (B) mismatch percentage from Illumina Induro-tRNA-seq or (C) total error percentage at position 31 from Nano-tRNA-seq. Bars show mean mismatch rate with SEM error bars; individual dots represent biological replicates (n = 3 per age group). Young (blue) and old (red) samples were compared by two-sided Welch’s t-test for each tRNA (***, p < 0.001; **, p < 0.01; *, p < 0.05; ns, not significant). (D) Modification co-occurrence network comparing young (left) and old (right) yeast samples. Nodes represent modification positions (9, 26, 32, 34, 37, 58) and aminoacylation (charging) status; node size is proportional to degree. Edges connect position pairs with significant co-occurrence (odds ratio, OR), assessed by equal-depth subsampling (300 reads per test, 200 iterations) with Benjamini-Hochberg correction (p < 0.05). Solid edges indicate co-occurrences consistently significant in >50% of iterations; dashed edges indicate borderline co-occurrences (25-50% of iterations). Edge width is proportional to |log2(OR)|; blue edges denote positive association (OR > 1), red edges denote negative association (OR < 1). Edge labels show mean OR across iterations.

tRNA modifications are hierarchically coordinated, with earlier modifications influencing downstream installation. To assess whether the altered m^1^A_58_ and m^3^C_32_ reflect a broader dysregulation, we applied the SLAC statistical framework for single-molecule analysis of modification crosstalk.^30^ This framework tests whether modification states at different positions co-occur on individual tRNA reads more or less often than expected by chance. For example, if a tRNA molecule carries m^1^A_58_ (detected as mismatch at position 58), does it also carry m^2^G/m^2^_2_G_26_ (mismatch at position 26) more often than a molecule without m^1^A_58_? To control for coverage difference across samples, we applied equal-depth subsampling (300 reads per test, 200 iterations) and retained only crosstalks consistently significant in >50% of iterations. After coverage correction, young yeast cells exhibit a robust co-occurrence network of five associations (Figure 5D). The core of this network is a positive triangle linking positions 58 (m^1^A_58_), 26 (m^2^G/m^2^_2_G) and 9 (m^1^G), with the strongest edge between positions 26 and 58 (OR = 6.6). Strikingly, this network is largely absent in aging cells (Figure 5D). The 26-58-9 triangle is not detected, consistent with age-associated disruption of coordinated modifications deposition. Notably, the global modification levels of m^1^G_9_ and m^2^G/m^2^_2_G_26_ remain unchanged with aging, indicating that the loss of co-occurrence reflects altered inter-position coordination on individual tRNA molecules rather than a uniform decrease in modification stoichiometry.

## DISCUSSION

Here we report a benchmarked Nanopore tRNA-sequencing resource that provides a comprehensive, single-molecule atlas of tRNAome remodeling during replicative aging in *S. cerevisiae*. This resource, validated with IVT controls and orthogonal Illumina sequencing, is available to the community for further exploration. More strikingly, we observed a global terminal cleavage at the 3’ CCA end of all mature tRNAs, correlating with increased proteostasis stress. We propose that Rny1 could be the enzyme that mediates this aging-associated cleavage. In addition, Nanopore sequencing detected significant aging-associated remodeling of modifications around the T loop on tRNA^Lys-CTT^, specifically involving m^5^U_54_, Ψ_55_ and m^1^A_58_. Beyond the T loop, we identified aging-associated changes at m^3^C sites in the anticodon loop, detected by both Nanopore and Illumina sequencing, revealing that modification remodeling during aging extends across multiple structural regions of the tRNA. Lastly, aging disrupts the interdependent deposition of modifications that normally ensures proper tRNA folding and function.

A novel mechanism to globally affect proteostasis is the terminal A cleavage at the 3’ CCA end. This cleavage showed a significant and global increase across tRNA species upon aging (Figure 3). One caveat is that the Nanopore splint adaptor ligation depends on an intact 3’ CCA end, so reduced terminal-A coverage could in principle reflect ligation bias or RNA degradation introduced during library preparation rather than *in vivo* CCA cleavage. Two observations argue against a purely preparation-derived artifact: young and old samples were processed in parallel under identical conditions, and the orthogonal Illumina charging protocol, which interrogates 3’ end integrity without splint ligation, independently shows a significant age-dependent loss of charged, CCA-intact tRNAs (Figure 3E). Nevertheless, absolute CCA-loss fractions should be interpreted as relative, platform-dependent estimates. Such cleavage would efficiently decrease the available pool of functional aminoacylated tRNAs to down-regulate protein synthesis as observed in aging.^31–33^ Previously, this process had only been observed under oxidative stress^26,34,35^ in both yeast and human cell line. In yeast and human cells, CCA-end cleavage has been observed under oxidative stress.^8,34^ It remains to be tested whether this is a general mechanism across other aging models. We propose Rny1 could be the RNase contributing to this cleavage based on genetic evidence, but *in vitro* biochemical validation will be needed in the future. While Rny1 has been shown to cleave tRNA anticodon loop upon oxidative stress,^36^ its specific role in aging has not been investigated. Further studies must determine if Rny1 requires uncharging to happen prior to the terminal A cleavage, and whether terminal A cleavage is linked to the anticodon loop cleavage.

The modifications m^5^U_54_, Ψ_55_ and m^1^A_58_ form a highly conserved T-loop circuit that critically stabilizes the tertiary structure of the tRNA.^17^ Each modification plays a distinct role in maintaining tRNA structural rigidity and functional activity. Recent work combining Nanopore direct RNA sequencing and mass spectrometry demonstrated that these modifications also function as an interconnected regulatory circuit, with Ψ_55_ promoting subsequent m^1^A_58_ modification on a substantial subset of yeast tRNAs. In particular, loss of Ψ_55_ resulted in reduced m^1^A_58_ modification across 19 of 42 cytosolic tRNA isoacceptors, indicating that modification at position 55 can influence methylation at the adjacent A58 position. Interestingly, this relationship was tRNA-specific: some tRNAs required Ψ_55_ for efficient m^1^A_58_ modification, whereas others retained m^1^A_58_ independently of Ψ_55_. The study further identified tRNA-Lys-CTT as one of several tRNAs in which m^1^A_58_ increased under nutrient-depleted conditions, demonstrating that T-loop modification status can also be dynamically regulated by cellular state.^17^ Alterations in this specific modification network can profoundly impact tRNA folding, aminoacylation efficiency, and subsequent interactions with the translation machinery. Strikingly, our data reveal a disruption of this circuit on tRNA^Lys-CTT^ (Figure 4C): while Ψ_55_ levels decreased, m^1^A_58_ simultaneously increased, the opposite of the canonical Ψ_55_ - m^1^A_58_ relationship. These findings suggest that aging might uncouple the interdependent modification circuit, potentially through altered activity or regulation of the respective modifying enzymes (Pus4 for Ψ_55_, Gcd10/Gcd14 for m^1^A_58_), rather than through the canonical sequential dependency.

Several recent reports have demonstrated polybasic (lysine and arginine) sequences are prone to ribosome stalling by slowed translation elongation during aging in *C.elegans*, *S. cerevisae*, and killifish species.^4,37^ Crucially, this stalling is highly specific to the AAG codon rather than AAA codon. An opportunity enabled by this resource is to test whether the specific changes observed in the tRNA^Lys-CTT^ T-loop affect translation of AAG-enriched transcripts, which encode proteins highly enriched in ribosome-associated pathways.^38^ The modification shift in tRNA^Lys-CTT^ (Figure 4) could lead to altered translation efficiency, presumably by affecting the interaction between the tRNA and eEF1A.^39,40^ The upstream cause of these specific changes in tRNA^Lys-CTT^ remains to be determined. Interestingly, tRNA^Lys-CTT^ is the only tRNA known to shuttle between the cytoplasm and mitochondria, a process mediated by lysine aminoacyltransferase MSK1.^41–44^ The AAG codon enrichment in ribosome-associated proteins together with the tRNA^Lys-CTT^ modification shifts we observe may point to a mechanism for maintaining ribosome biosynthesis during aging.

Lastly, our Nanopore sequencing pipeline provides a robust resource for the systematic interrogation of tRNA expression and modification patterns. By analyzing base-calling errors and comparing them with unmodified IVT tRNAs, we demonstrated that 87.2% of annotated modifications can be detected (Figure 1), with the modification type-specific detection rates characterized across both platforms (Figure 2). The remaining 12.8% will need more careful evaluation. For example, m^3^C fell below the Nanopore error-based detection threshold at the modification sites. By using error signature at the −1 position, we were able to identify age-specific m^3^C from Nanopore, which is validated by Illumina platform (Figure 5). It is possible that modification-aware basecaller will further improve detection of these modification types in the future.^18^ Notably, the systematic comparison provides a starting point for the discovery of novel tRNA modifications, especially for the sites that are above error threshold and not adjacent to other modifications (Figure 1D). We provide the complete Nanopore and Illumina output as Data S3. We anticipate that this resource will serve as a foundation for systematic investigations of tRNA dynamics not only in aging but across diverse biological contexts.

## RESOURCE AVAILABILITY

### Lead contact

Requests for further information and resources should be directed to and will be fulfilled by the lead contact, Zhangli Su.

### Materials availability

This study did not generate new unique reagents.

### Data and code availability

Nanopore and Illumina data has been deposited to Gene Expression Omnibus under accession numbers GSE346842 and GSE346844.

Any additional information required to reanalyze the data reported in this paper is available from the lead contact upon request.

## ACKNOWLEDGMENTS

We would like to thank Shane Rich-New and IT Research Computing Team at the University of Alabama at Birmingham for NGS support. This research was supported by NIH grants R00CA259526 (to Z.S.), and T32GM135028 (to K.M.B.). This work was also supported by NIH grants R01GM075240, R21AG08588, R01AG089953 (to J.S.S.) and F31AG081044 (to L.N.P.).

## AUTHOR CONTRIBUTIONS

Conceptualization, Z.S. and J.S.; methodology, data curation, formal analysis, visualization, K.M.B. and Z.S.; validation, C.S. and Z.S.; resources, C.P., N.Z., L.N.P.; writing - original draft, K.M.B. and Z.S.; writing - review & editing, C.P., N.Z., L.N.P., C.S. and J.S.; supervision, project administration, funding acquisition, Z.S. and J.S.

## SUPPLEMENTAL INFORMATION

Figures S1-S5

Data S1-S3

## METHODS

### EXPERIMENTAL MODEL DETAILS

Replicative aging was performed using the Mini Chemostat Aging Device (MAD) system similar to previously.^24^ Briefly, log-phase BY4741 cells (OD_600_ of ∼1.2-1.7) were biotinylated with EZ-Link Sulfo-NHS- LC-Biotin (Thermo Scientific) and bound to magnetic streptavidin beads (Dynabeads MyOne Streptavidin C, Invitrogen). Unbiotinylated daughter cells were removed and retained as the young control. Bead-bound mother cells were immobilized in chemostat tubes with neodymium magnets and aged for 36 hours (∼8 generations) with continuous perfusion of fresh SC medium. After aging, mother cells were harvested and purified from remaining daughter cells using magnetic racks. Replicative age was verified by calcofluor white staining and imaging at 60X on an EVOS M7000 microscope (Thermo Scientific). The *RNY1* overexpression strain is derived from the MoBY 2.0 collection (PMID: 19349972), consisting of individual yeast genes inserted into the 2µ high copy vector p5476 and under control of its endogenous promoter. The *GCD14* knockdown strain is derived from the DAmP collection (PMID: 18622397), consisting of essential genes with a kanMX4 selectable marker inserted into the 3’-UTR, yielding a hypomorphic allele.

### METHOD DETAILS

#### tRNA isolation

Total RNA was extracted from 150 million cells using standard hot acidic phenol method.^45^ To enrich for tRNAs, total RNA was resolved on a denaturing polyacrylamide gel (10% TBE-Urea gel). The gel region corresponding to ∼70 - 90 nt was excised, and tRNAs were eluted in 0.3 M NaCl overnight at 4°C, followed by ethanol precipitation. RNA concentration was assessed using a Nanodrop spectrophotometer.

#### *In vitro* transcription of tRNAs

Oligo pools of 54 mature tRNA sequences were ordered from IDT and included a T7 promoter. The oligo pool was dissolved in low EDTA TE buffer to a concentration of 100 ng/ml. Templates for *in vitro* transcription were generated by 3-step PCR amplification with NEB Q5 Polymerase. PCR product size was verified by agarose gel. The remainder of PCR product was purified using DNA Clean and Concentrator Kit (Zymo, cat #D4014). The purified products were digested with BsaI-HFv2 (NEB, cat #R3733S) to ensure correct termination at CCA end. Digestion products were purified with DNA Clean and Concentrator kit (Zymo, cat #D4014). *In vitro* transcription was performed using HiScribe T7 High Yield RNA synthesis kit (NEB, cat #E2040S) at 37°C for 3 hours. The transcription reaction contained 7.5 mM of each NTP (ATP, CTP, UTP, GTP), 10X reaction buffer (40 mM Tris-HCl pH 7.9, 6 mM MgCl2, 2 mM spermidine, 1 mM DTT, 50 units/ml T7 RNA polymerase, 5 mM DTT, RNase Inhibitor (Life Technologies, cat #AM2696), and 2.5% PEG8000. The transcript products were treated with DNase I for 15 minutes at 37°C. RNA products were then cleaned by RNA Clean and Concentrator kit (Zymo, cat #R1016). To convert 5’ triphosphate into 5’ monophosphate, RNA products were treated with 1 μl RppH per 100 ng of RNA with 1x NEBuffer 2 (NEB, cat #M0356S), and RNase Inhibitor at 37°C for 2 hours. The reaction was inactivated by adding 0.6 μl of 500 mM EDTA and incubating at 65°C for 5 minutes and left to cool slowly to room temperature. The RppH digested product was purified by 10% Novex Urea denaturing polyacrylamide gel electrophoresis (Fisher Scientific, cat #EC68752BOX), size selected for digested product and purified by ethanol precipitation.

#### Nanopore library preparation

Gel-purified tRNAs were prepared for sequencing using the Nano-tRNA-Seq Kit (Immagina Biotechnologies, cat #NTRSQ-12) according to manufacturer’s instructions with the following modifications: tRNAs were purified by 10% Novex Urea denaturing polyacrylamide gel electrophoresis (Fisher Scientific, cat #EC68752BOX) instead of the kit’s small RNA enrichment method. Briefly, the protocol includes adaptor ligation, barcoding for multiplexing, and purification steps optimized for small RNAs. Libraries were quantified by Qubit HS dsDNA assay (Fisher Scientific, cat #Q32851) and loaded onto an Oxford Nanopore MinION RNA flow cell (cat #FLO-MIN004RA) and sequencing was performed following the kit guidelines with MinION Mk1B.

#### Nano-tRNA-seq data analysis

Raw Nanopore reads were processed using the Nano-tRNA-Seq kit guidelines, including basecalling with Dorado version 1.0.2, mapping to reference tRNA sequences using minimap2 version 2.30, and demultiplexing. tRNA reference dataset was downloaded from https://github.com/novoalab/Nano-tRNAseq/ and counts were quantified using the get_counts.py script.^8^ Differential expression analysis between experimental groups was performed using DESeq2.^46^ CCA end integrity was assessed by by measuring per-read coverage at the three terminal positions (C1, C2, A) of the 3’ CCA-end. Normalized CCA coverage was calculated as the ratio of coverage at each terminal position relative to the first C position (C1), yielding norm_C2 = C2_coverage / C1_coverage and norm_A = A_coverage / C1_coverage. This normalization accounts for differences in overall tRNA abundance, enabling comparison of CCA-end completeness across samples and age groups.

Basecalling error statistics were measured using EpiNano_Variants.py version 1.2, and total error rate was calculated by summing mismatch, insertion, and deletion error rates.^47^ To establish a modification-free baseline for threshold calibration, all 54 unique mature tRNA sequences were *in vitro* transcribed (IVT) lacking endogenous modifications and sequenced alongside biological samples. Receiver-Operator Characteristic (ROC) analysis was performed using the pROC package^48^ in R version 4.4.3, with true positives defined as positions annotated by MODOMICS^19^ on the 31 annotated tRNAs and true negatives defined as all positions from the IVT dataset. Error rates were averaged across all six biological replicates prior to ROC analysis. To determine the optimal error metric, ROC curves were computed for each error component individually (mismatch, insertion, deletion) and for total error (mismatch + insertion + deletion). To account for the neighbor effect whereby Nanopore base-calling errors propagate to positions adjacent to modified bases, error rates were smoothed using a ± k-nucleotide sliding window (k = 1, 2, 3), taking the maximum total error rate within each window. The optimal classification threshold was selected using Youden’s J statistic (maximizing sensitivity + specificity − 1).

For metagene analysis of age-dependent error changes (Figure 3A), smoothing was performed using the mean error change within a ± 3-nucleotide window. Differential modification analysis between experimental groups was performed using Student’s t-test in R version 4.4.3, and results were visualized in heatmaps and other summary boxplots.

#### Illumina library preparation

Total RNAs were first processed by peroxide oxidation, beta-elimination, and deacylation^49,50^ to remove aminoacylated 3’ CCA ends and generate uniform 3’ hydroxyl termini. Briefly, RNAs were oxidized with 150 mM sodium periodate (NaIO_4_) for 30 min at room temperature, quenched with 1 M glucose for 5 min, then subjected to beta-elimination and deacylation with 100 mM sodium borate and SUPERaseIn at 37°C for 30 min. The 3’ ends were repaired with T4 PNK (NEB) for 30 min at 37°C, followed by heat inactivation at 65°C for 10 min. Small RNA fractions (<200 nt) were enriched using the Zymo RNA Clean and Concentrator kit with two columns per sample. Libraries were prepared from 25 ng of processed small RNAs using the NEBNext Low Bias Library Prep Kit (NEB) with the following modifications: reverse transcription was performed using Induro RT enzyme (NEB) with an extended incubation of overnight at 42°C to maximize read-through of tRNA modifications.^50^ Post-RT cleanup was performed using 0.5X SparQ beads (QuantBio) with 10 μL isopropanol instead of the manufacturer’s size selection protocol, as small RNAs had been pre- enriched by column purification. Libraries were amplified by PCR for 13 cycles using NEBNext LV Unique Dual Index Primers. For RNY1 over-expression samples, total RNA was treated with T4 PNK with 1 mM ATP at 37 °C for 1 hour to convert 3’ cP (or 3’ P) and 5’ OH termini to 3’ OH and 5’ P ends. Treated RNA was cleaned up prior to library preparation with NEBNext Low Bias Library Prep Kit (NEB).

#### Illumina library data analysis and modification co-occurrence analysis

Briefly, reads were trimmed with cutadapt, and reads longer than 50 nucleotides long are aligned to the *S. cerevisiae* tRNA reference using mim-tRNA-seq^16,49^ with the crosstalk analysis setting. mim-tRNA-seq quantifies tRNA abundance and detects base modifications via reverse transcription (RT) misincorporation signatures at each canonical position. Differential expression analysis between young and old samples was performed using DESeq2^46^ on mim-tRNA-seq count output. For modification analysis, RT misincorporation proportions were extracted per isodecoder and canonical position from the mim-tRNA-seq mismatch table. Differential modification analysis between age groups was performed using Student’s t-test in R version 4.4.3 on misincorporation proportions at known modification positions, with multiple testing correction applied where appropriate.

Per-read modification co-occurrence was computed using the mim-tRNA-seq crosstalk module^30^, which evaluates pairwise co-occurrence of RT misincorporation signatures across canonical positions on individual reads. For each pair of positions (var1, var2) within each tRNA and sample, a 2×2 contingency table was constructed counting reads with misincorporation at both positions (True/True), at one position only (True/False, False/True), or at neither (False/False). The odds ratio (OR) was calculated as (True/True × False/False) / (True/False × False/True), and statistical significance was assessed using Fisher’s exact test, with Benjamini-Hochberg correction for multiple testing. Co-occurrence networks were visualized as undirected graphs using igraph^51^ in R, where nodes represent canonical modification positions (e.g., positions 9, 26, 32, 34, 37, 58) or charging status ("Charged"), and edges represent significant co- occurrence pairs (corrected p < 0.05). Edge width was proportional to the absolute log₂(OR) (effect size), and edge color indicated direction: positive associations (OR > 1, co-occurrence exceeding expectation) in blue and negative associations (OR < 1, mutual exclusivity) in red. Networks were constructed separately for young and old samples. To control for read depth differences between age groups, a subsampling analysis was performed in which contingency tables were repeatedly subsampled to matched read depths, and Fisher’s exact test was recomputed across iterations to assess robustness of significant co- occurrences. Co-occurrence edges significant in >50% of subsampling iterations were classified as consistent (solid lines), while edges significant in 25-50% of iterations were retained as near-consistent context (dashed lines).

### QUANTIFICATION AND STATISTICAL ANALYSIS

The statistical information of each experiment, including the statistical method, the p value and sample numbers (n) are shown in corresponding figure legends and methods.

## REFERENCES

1. López-Otín, C., Blasco, M.A., Partridge, L., Serrano, M., and Kroemer, G. (2023). Hallmarks of aging: An expanding universe. Cell 186, 243–278. 10.1016/j.cell.2022.11.001.

2. Hipp, M.S., Kasturi, P., and Hartl, F.U. (2019). The proteostasis network and its decline in ageing. Nat Rev Mol Cell Biol 20, 421–435. 10.1038/s41580-019-0101-y.

3. Walther, D.M., Kasturi, P., Zheng, M., Pinkert, S., Vecchi, G., Ciryam, P., Morimoto, R.I., Dobson, C.M., Vendruscolo, M., Mann, M., et al. (2015). Widespread Proteome Remodeling and Aggregation in Aging C. elegans. Cell 161, 919–932. 10.1016/j.cell.2015.03.032.

4. Stein, K.C., Morales-Polanco, F., van der Lienden, J., Rainbolt, T.K., and Frydman, J. (2022). Ageing exacerbates ribosome pausing to disrupt cotranslational proteostasis. Nature 601, 637–642. 10.1038/s41586-021-04295-4.

5. Woodward, K., and Shirokikh, N.E. (2021). Translational control in cell ageing: an update. Biochem Soc Trans 49, 2853–2869. 10.1042/BST20210844.

6. Hu, Z., Xia, B., Postnikoff, S.D., Shen, Z.-J., Tomoiaga, A.S., Harkness, T.A., Seol, J.H., Li, W., Chen, K., and Tyler, J.K. (2018). Ssd1 and Gcn2 suppress global translation efficiency in replicatively aged yeast while their activation extends lifespan. Elife 7, e35551. 10.7554/eLife.35551.

7. Gerashchenko, M.V., Peterfi, Z., Yim, S.H., and Gladyshev, V.N. (2021). Translation elongation rate varies among organs and decreases with age. Nucleic Acids Res 49, e9. 10.1093/nar/gkaa1103.

8. Lucas, M.C., Pryszcz, L.P., Medina, R., Milenkovic, I., Camacho, N., Marchand, V., Motorin, Y., Ribas de Pouplana, L., and Novoa, E.M. (2024). Quantitative analysis of tRNA abundance and modifications by nanopore RNA sequencing. Nat Biotechnol 42, 72–86. 10.1038/s41587-023-01743-6.

9. White, L.K., Dobson, K., Del Pozo, S., Bilodeaux, J.M., Andersen, S.E., Baldwin, A., Barrington, C., Körtel, N., Martinez-Seidel, F., Strugar, S.M., et al. (2024). Comparative analysis of 43 distinct RNA modifications by nanopore tRNA sequencing. Preprint, 10.1101/2024.07.23.604651 https://doi.org/10.1101/2024.07.23.604651.

10. Thomas, N.K., Poodari, V.C., Jain, M., Olsen, H.E., Akeson, M., and Abu-Shumays, R.L. (2021). Direct Nanopore Sequencing of Individual Full Length tRNA Strands. ACS Nano 15, 16642–16653. 10.1021/acsnano.1c06488.

11. Filer, D., Thompson, M.A., Takhaveev, V., Dobson, A.J., Kotronaki, I., Green, J.W.M., Heinemann, M., Tullet, J.M.A., and Alic, N. (2017). RNA polymerase III limits longevity downstream of TORC1. Nature 552, 263–267. 10.1038/nature25007.

12. Malik, Y., Kulaberoglu, Y., Anver, S., Javidnia, S., Borland, G., Rivera, R., Cranwell, S., Medelbekova, D., Svermova, T., Thomson, J., et al. (2024). Disruption of tRNA biogenesis enhances proteostatic resilience, improves later-life health, and promotes longevity. PLoS Biol 22, e3002853. 10.1371/journal.pbio.3002853.

13. McCormick, M.A., Delaney, J.R., Tsuchiya, M., Tsuchiyama, S., Shemorry, A., Sim, S., Chou, A.C.- Z., Ahmed, U., Carr, D., Murakami, C.J., et al. (2015). A Comprehensive Analysis of Replicative Lifespan in 4,698 Single-Gene Deletion Strains Uncovers Conserved Mechanisms of Aging. Cell Metab 22, 895–906. 10.1016/j.cmet.2015.09.008.

14. Gong, R.-Z., Yan, T.-M., Pan, Y., Cao, K.-Y., Cheng, Y.-T., Mo, L.-Y., and Jiang, Z.-H. (2026). Evolutionarily Conserved Decline of tRNA Mannosyl-Queuosine Links Translational Regulation to Aging and Is Reversed by Queuine. Preprint at bioRxiv, 10.64898/2026.03.22.713446 https://doi.org/10.64898/2026.03.22.713446.

15. Fu, Y., Jiang, F., Zhang, X., Pan, Y., Xu, R., Liang, X., Wu, X., Li, X., Lin, K., Shi, R., et al. (2024). Perturbation of METTL1-mediated tRNA N7- methylguanosine modification induces senescence and aging. Nat Commun 15, 5713. 10.1038/s41467-024-49796-8.

16. Behrens, A., Rodschinka, G., and Nedialkova, D.D. (2021). High-resolution quantitative profiling of tRNA abundance and modification status in eukaryotes by mim-tRNAseq. Molecular Cell 81, 1802–1815.e7. 10.1016/j.molcel.2021.01.028.

17. Shaw, E.A., Thomas, N.K., Jones, J.D., Abu-Shumays, R.L., Vaaler, A.L., Akeson, M., Koutmou, K.S., Jain, M., and Garcia, D.M. (2024). Combining Nanopore direct RNA sequencing with genetics and mass spectrometry for analysis of T-loop base modifications across 42 yeast tRNA isoacceptors. Nucleic Acids Res 52, 12074–12092. 10.1093/nar/gkae796.

18. Diensthuber, G., Pryszcz, L.P., Llovera, L., Lucas, M.C., Delgado-Tejedor, A., Cruciani, S., Roignant, J.-Y., Begik, O., and Novoa, E.M. (2024). Enhanced detection of RNA modifications and read mapping with high-accuracy nanopore RNA basecalling models. Genome Res 34, 1865–1877. 10.1101/gr.278849.123.

19. Cappannini, A., Ray, A., Purta, E., Mukherjee, S., Boccaletto, P., Moafinejad, S.N., Lechner, A., Barchet, C., Klaholz, B.P., Stefaniak, F., et al. (2024). MODOMICS: a database of RNA modifications and related information. 2023 update. Nucleic Acids Res 52, D239–D244. 10.1093/nar/gkad1083.

20. Bohn, P., Gribling-Burrer, A.-S., Ambi, U.B., and Smyth, R.P. (2023). Nano-DMS-MaP allows isoform-specific RNA structure determination. Nat Methods 20, 849–859. 10.1038/s41592-023-01862-7.

21. Kang, X., Zhang, K., Goyon, A., and Stephenson, W. (2026). Systematic assessment of diverse RNA modifications using nanopore direct RNA sequencing. Nucleic Acids Res 54, gkag411. 10.1093/nar/gkag411.

22. O’Laughlin, R., Jin, M., Li, Y., Pillus, L., Tsimring, L.S., Hasty, J., and Hao, N. (2020). Advances in quantitative biology methods for studying replicative aging in *Saccharomyces cerevisiae*. Translational Medicine of Aging 4, 151–160. 10.1016/j.tma.2019.09.002.

23. Longo, V.D., Shadel, G.S., Kaeberlein, M., and Kennedy, B. (2012). Replicative and Chronological Aging in Saccharomyces cerevisiae. Cell Metab 16, 18–31. 10.1016/j.cmet.2012.06.002.

24. Power, L.N., Zawrotna, N., Dinda, M., Weir, A.E., Paudel, B.B., Ghimire, O., Kisiel, K., Letai, C.T., Janes, K.A., and Smith, J.S. (2026). Age-dependent topoisomerase I depletion alters recruitment of rDNA silencing complexes. Journal of Biological Chemistry 302, 111062. 10.1016/j.jbc.2025.111062.

25. Hou, Y.-M. (2010). CCA Addition to tRNA: Implications for tRNA Quality Control. IUBMB Life 62, 251–260. 10.1002/iub.301.

26. Czech, A., Wende, S., Mörl, M., Pan, T., and Ignatova, Z. (2013). Reversible and Rapid Transfer- RNA Deactivation as a Mechanism of Translational Repression in Stress. PLOS Genetics 9, e1003767. 10.1371/journal.pgen.1003767.

27. Wang, L., and Lin, S. (2023). Emerging functions of tRNA modifications in mRNA translation and diseases. Journal of Genetics and Genomics 50, 223–232. 10.1016/j.jgg.2022.10.002.

28. Schultz, S.K., Katanski, C.D., Halucha, M., Peña, N., Fahlman, R.P., Pan, T., and Kothe, U. (2024). Modifications in the T arm of tRNA globally determine tRNA maturation, function, and cellular fitness. Proceedings of the National Academy of Sciences 121, e2401154121. 10.1073/pnas.2401154121.

29. Agris, P.F., Narendran, A., Sarachan, K., Väre, V.Y.P., and Eruysal, E. (2017). The Role of RNA Modifications in Translational Fidelity. Enzymes 41, 1–50. 10.1016/bs.enz.2017.03.005.

30. Hernandez-Alias, X., Katanski, C.D., Zhang, W., Assari, M., Watkins, C.P., Schaefer, M.H., Serrano, L., and Pan, T. (2023). Single-read tRNA-seq analysis reveals coordination of tRNA modification and aminoacylation and fragmentation. Nucleic Acids Res 51, e17. 10.1093/nar/gkac1185.

31. Zhou, Z., Sun, B., Yu, D., and Bian, M. (2021). Roles of tRNA metabolism in aging and lifespan. Cell Death Dis 12, 548. 10.1038/s41419-021-03838-x.

32. Solyga, M., Majumdar, A., and Besse, F. (2024). Regulating translation in aging: from global to gene- specific mechanisms. EMBO Rep 25, 5265–5276. 10.1038/s44319-024-00315-2.

33. Skariah, G., and Todd, P.K. (2021). TRANSLATIONAL CONTROL IN AGING AND NEURODEGENERATION. Wiley Interdiscip Rev RNA 12, e1628. 10.1002/wrna.1628.

34. Akiyama, Y., Lyons, S.M., Fay, M.M., Tomioka, Y., Abe, T., Anderson, P.J., and Ivanov, P. (2022). Selective Cleavage at CCA Ends and Anticodon Loops of tRNAs by Stress-Induced RNases. Front. Mol. Biosci. 9. 10.3389/fmolb.2022.791094.

35. Czech, A. (2020). Deep sequencing of tRNA’s 3’-termini sheds light on CCA-tail integrity and maturation. RNA 26, 199–208. 10.1261/rna.072330.119.

36. Thompson, D.M., and Parker, R. (2009). The RNase Rny1p cleaves tRNAs and promotes cell death during oxidative stress in Saccharomyces cerevisiae. J Cell Biol 185, 43–50. 10.1083/jcb.200811119.

37. Di Fraia, D., Marino, A., Lee, J.H., Kelmer Sacramento, E., Baumgart, M., Bagnoli, S., Balla, T., Schalk, F., Kamrad, S., Guan, R., et al. (2025). Altered translation elongation contributes to key hallmarks of aging in the killifish brain. Science 389, eadk3079. 10.1126/science.adk3079.

38. Ghanegolmohammadi, F., Ohnuki, S., Byrne, S., Raman, R., Begley, T.J., and Dedon, P.C. (2026). Synonymous codon usage defines functional gene families. BMC Biol 24, 44. 10.1186/s12915-026-02505-x.

39. He, R., Lv, Z., Li, Y., Ren, S., Cao, J., Zhu, J., Zhang, X., Wu, H., Wan, L., Tang, J., et al. (2024). tRNA-m1A methylation controls the infection of *Magnaporthe oryzae* by supporting ergosterol biosynthesis. Developmental Cell 59, 2931–2946.e7. 10.1016/j.devcel.2024.08.002.

40. Liu, Y., Zhou, J., Li, X., Zhang, X., Shi, J., Wang, X., Li, H., Miao, S., Chen, H., He, X., et al. (2022). tRNA-m1A modification promotes T cell expansion via efficient MYC protein synthesis. Nat Immunol 23, 1433–1444. 10.1038/s41590-022-01301-3.

41. Tarassov, I., Entelis, N., and Martin, R.P. (1995). Mitochondrial import of a cytoplasmic lysine-tRNA in yeast is mediated by cooperation of cytoplasmic and mitochondrial lysyl-tRNA synthetases. The EMBO Journal 14, 3461–3471. 10.1002/j.1460-2075.1995.tb07352.x.

42. Entelis, N.S., Kieffer, S., Kolesnikova, O.A., Martin, R.P., and Tarassov, I.A. (1998). Structural requirements of tRNALys for its import into yeast mitochondria. Proceedings of the National Academy of Sciences 95, 2838–2843. 10.1073/pnas.95.6.2838.

43. Kamenski, P., Kolesnikova, O., Jubenot, V., Entelis, N., Krasheninnikov, I.A., Martin, R.P., and Tarassov, I. (2007). Evidence for an adaptation mechanism of mitochondrial translation via tRNA import from the cytosol. Mol Cell 26, 625–637. 10.1016/j.molcel.2007.04.019.

44. Kolesnikova, O., Kazakova, H., Comte, C., Steinberg, S., Kamenski, P., Martin, R.P., Tarassov, I., and Entelis, N. (2010). Selection of RNA aptamers imported into yeast and human mitochondria. RNA 16, 926–941. 10.1261/rna.1914110.

45. Collart, M.A., and Oliviero, S. (1993). Preparation of Yeast RNA. Current Protocols in Molecular Biology 23, 13.12.1–13.12.5. 10.1002/0471142727.mb1312s23.

46. Love, M.I., Huber, W., and Anders, S. (2014). Moderated estimation of fold change and dispersion for RNA-seq data with DESeq2. Genome Biol 15, 550. 10.1186/s13059-014-0550-8.

47. Liu, H., Begik, O., Lucas, M.C., Ramirez, J.M., Mason, C.E., Wiener, D., Schwartz, S., Mattick, J.S., Smith, M.A., and Novoa, E.M. (2019). Accurate detection of m6A RNA modifications in native RNA sequences. Nat Commun 10, 4079. 10.1038/s41467-019-11713-9.

48. Robin, X., Turck, N., Hainard, A., Tiberti, N., Lisacek, F., Sanchez, J.-C., and Müller, M. (2011). pROC: an open-source package for R and S+ to analyze and compare ROC curves. BMC Bioinformatics 12, 77. 10.1186/1471-2105-12-77.

49. Behrens, A., and Nedialkova, D.D. (2022). Experimental and computational workflow for the analysis of tRNA pools from eukaryotic cells by mim-tRNAseq. STAR Protocols 3, 101579. 10.1016/j.xpro.2022.101579.

50. Nakano, Y., Gamper, H., McGuigan, H., Maharjan, S., Li, J., Sun, Z., Yigit, E., Grünberg, S., Krishnan, K., Li, N.-S., et al. (2025). Genome-wide profiling of tRNA modifications by Induro-tRNAseq reveals coordinated changes. Nat Commun 16, 1047. 10.1038/s41467-025-56348-1.

51. Antonov, M., Csárdi, G., Horvát, S., Müller, K., Nepusz, T., Noom, D., Salmon, M., Traag, V., Welles, B.F., and Zanini, F. (2023). igraph enables fast and robust network analysis across programming languages. Preprint at arXiv, 10.48550/ARXIV.2311.10260 https://doi.org/10.48550/ARXIV.2311.10260.

