## Supplementary Figures for "Integrative Nanopore and Illumina sequencing reveals age-associated tRNA modification and CCA-tail dynamics in yeast"

#### SUPPLEMENTARY FIGURE TITLES AND LEGENDS

##### Figure S1. Global profiling of tRNA expression and modifications using Nano-tRNA-seq.

(A) $Log_2$ comparison of tRNA counts between the present study and *Lucas et al.*
(B) Urea gel showing the size and digest of *in vitro* transcribed (IVT) tRNAs used in this study.
(C) Correlation of total error rates at $m^1A$ sites between the present study and *Lucas et al.*
(D) Correlation of total error rates at $m^1A$ sites between the present study and *Behrens et al.*
(E) 10% TBU-Urea PAGE gel of *in vitro* transcribed library.
(F-G) ROC curves comparing individual basecalling error signatures (F) or total error with smoothing windows (G) for classifying MODOMICS-annotated modification sites (true positives, n = 375) against IVT positions (true negatives, n = 3,252). For each window size, the smoothed error rate at a given position was defined as the maximum total error within the ± k-nucleotide window.

##### Figure S2. Nano-tRNA-seq reveals modification-specific error signatures in yeast tRNAs.

(A) Error type composition (mismatch, insertion, deletion) for 5 additional modification types. Format as in Figure 2A; bars show mean proportion of each error type within ±2 positions of modification sites, averaged across all instances per modification type.
(B-C) Heatmaps showing position-specific mismatch rates across ±5 positions surrounding annotated modification sites, for Nanopore (B) and Illumina (C). Each row represents a modification type; columns represent sequential positions relative to the modification site (0). Color scale spans 0 to the maximum observed mismatch rate per platform.

##### Figure S3. tRNA expression is stable during aging while CCA tails undergo cleavage.

(A) Integrated Genome Viewer (IGV) screenshots illustrating CCA tail coverage for representative tRNAs.
(B-D) C-ending (B), CCA-ending (C) and CC-ending (D) fractions in RNY1 OE versus control by T4 PNK Illumina analysis. Bars show mean across cytoplasmic tRNAs with individual values overlaid. Statistics by paired t-test with Bonferroni correction (*** $p < 0.001$, NS $p > 0.05$).

##### Figure S4. Replicative aging alters specific tRNA T-loop modification signatures.

(A) Heatmap showing the percent change in error rate between old and young samples with Nanopore sequencing. Positive values (red) indicate higher error in old samples, negative values (blue) indicate higher error in young samples.
(B) Representative IGV screenshots showing mismatch patterns at specific modification sites in Lys-CTT tRNAs in young and old samples.
(C) Alignment of Lys-TTT and Lys-CTT mature tRNA sequences. Bold letters indicate nucleotide positions that differ between the two sequences, and underlined letters denote the anticodon.
(D) Bar plots show mean Nanopore error rate (± SEM, n = 3 biological replicates) for $tRNA^{Lys\text{-}TTT}$ at positions 54 ($m^5U$), 55 (Ψ), and 58 ($m^1A$). Blue bars = young, red bars = old; black dots = individual replicate values. Significance by two-sided Student's t-test: * $p < 0.05$, ** $p < 0.01$, *** $p < 0.001$.
(E) MA plot of age-dependent mismatch changes at position 58 ($m^1A$). Each point represents one of 42 cytoplasmic tRNAs; x-axis shows the Illumina mismatch rate in young (%), y-axis shows the change (old − young, %). Grey points = not significant; red = significant increase in old, blue = significant decrease (two-sided Student's t-test, $p < 0.05$). Significant tRNAs are labeled.

##### Figure S5. Age-dependent $m^3C_{32}$ increase and altered modification co-occurrence.

(A) Mismatch percentage with Illumina sequencing by canonical positions. Each dot represents one tRNA.

(B) Heatmap showing the percent change in error rate between old and young samples with Illumina sequencing. Positive values (red) indicate higher error in old samples, negative values (blue) indicate higher error in young samples.

### Figure S1

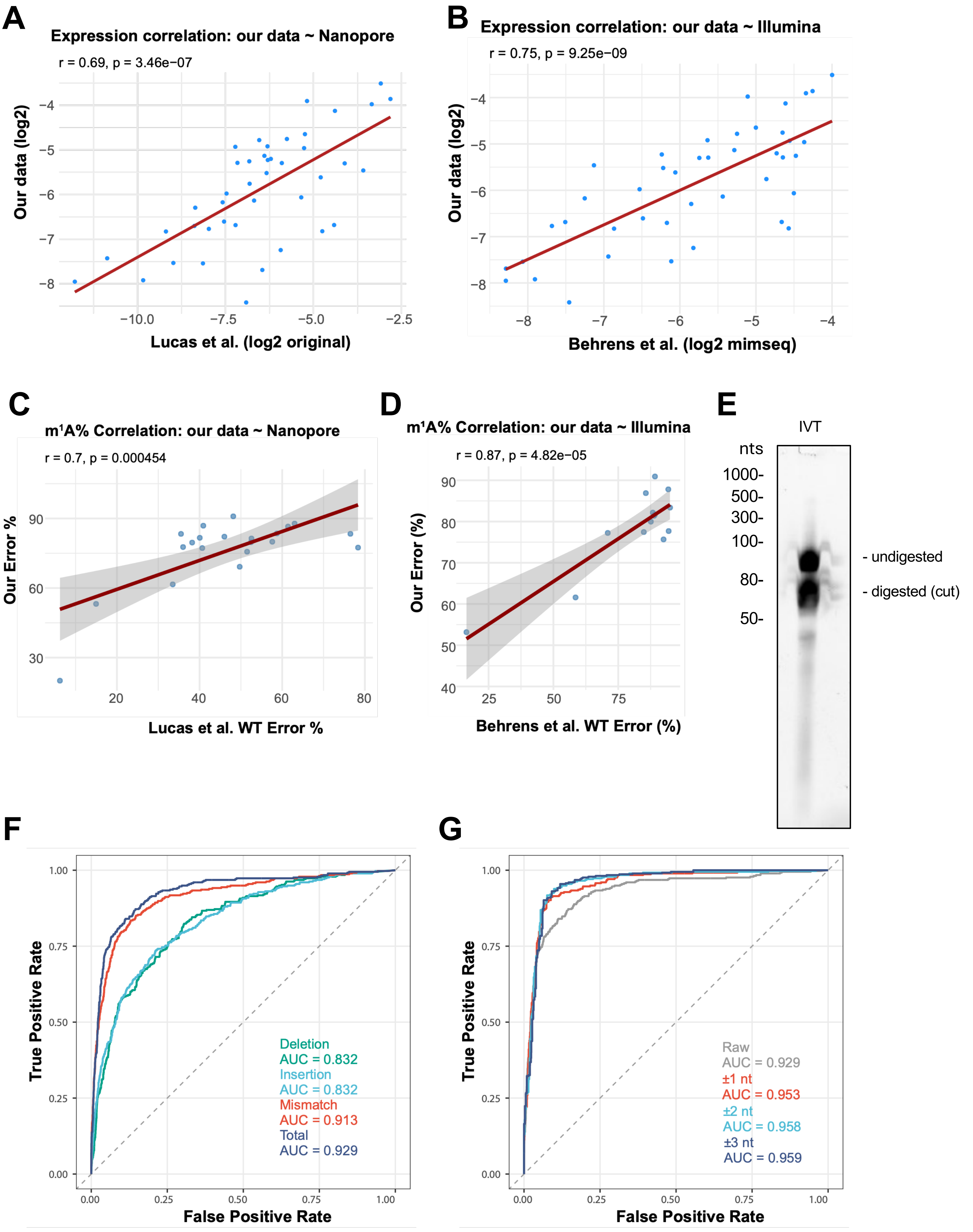

A
Expression correlation: our data ~ Nanopore
r = 0.69, p = 3.46e−07
Our data (log2)
Lucas et al. (log2 original)
B
Expression correlation: our data ~ Illumina
r = 0.75, p = 9.25e−09
Our data (log2)
Behrens et al. (log2 mimseq)
C
m¹A% Correlation: our data ~ Nanopore
r = 0.7, p = 0.000454
Our Error %
Lucas et al. WT Error %
D
m¹A% Correlation: our data ~ Illumina
r = 0.87, p = 4.82e−05
Our Error (%)
Behrens et al. WT Error (%)
E
IVT
nts
1000-
500-
300-
100-
80-
50-
- undigested
- digested (cut)
F
True Positive Rate
False Positive Rate
Deletion
AUC = 0.832
Insertion
AUC = 0.832
Mismatch
AUC = 0.913
Total
AUC = 0.929
G
True Positive Rate
False Positive Rate
Raw
AUC = 0.929
±1 nt
AUC = 0.953
±2 nt
AUC = 0.958
±3 nt
AUC = 0.959

### Figure S2

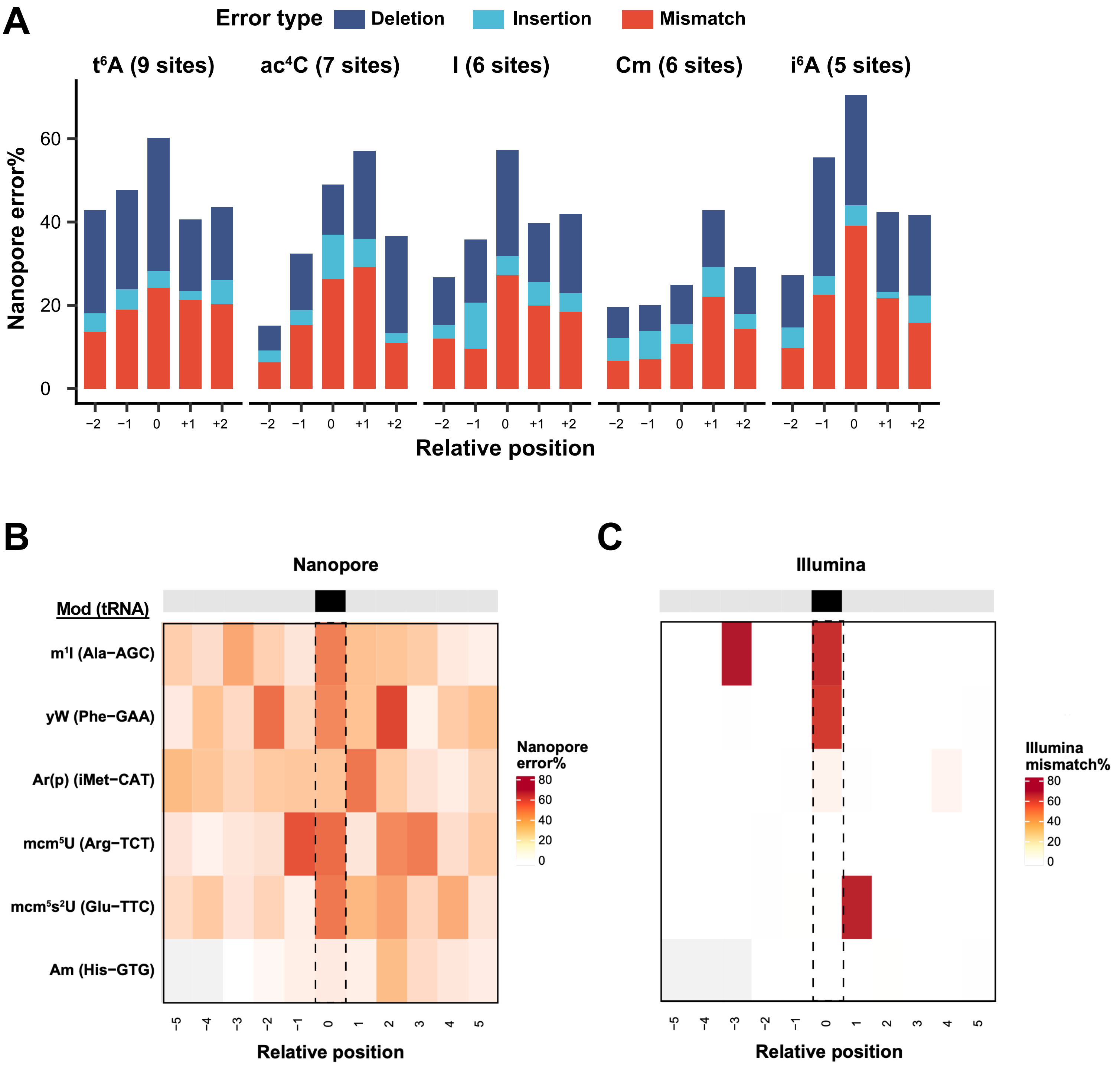

A
Error type
Deletion
Insertion
Mismatch
t6A (9 sites)
ac4C (7 sites)
I (6 sites)
Cm (6 sites)
i6A (5 sites)
Nanopore error%
Relative position
B
Nanopore
Mod (tRNA)
m1I (Ala−AGC)
yW (Phe−GAA)
Ar(p) (iMet−CAT)
mcm5U (Arg−TCT)
mcm5s2U (Glu−TTC)
Am (His−GTG)
Nanopore error%
Relative position
C
Illumina
Illumina mismatch%
Relative position

Figure S3

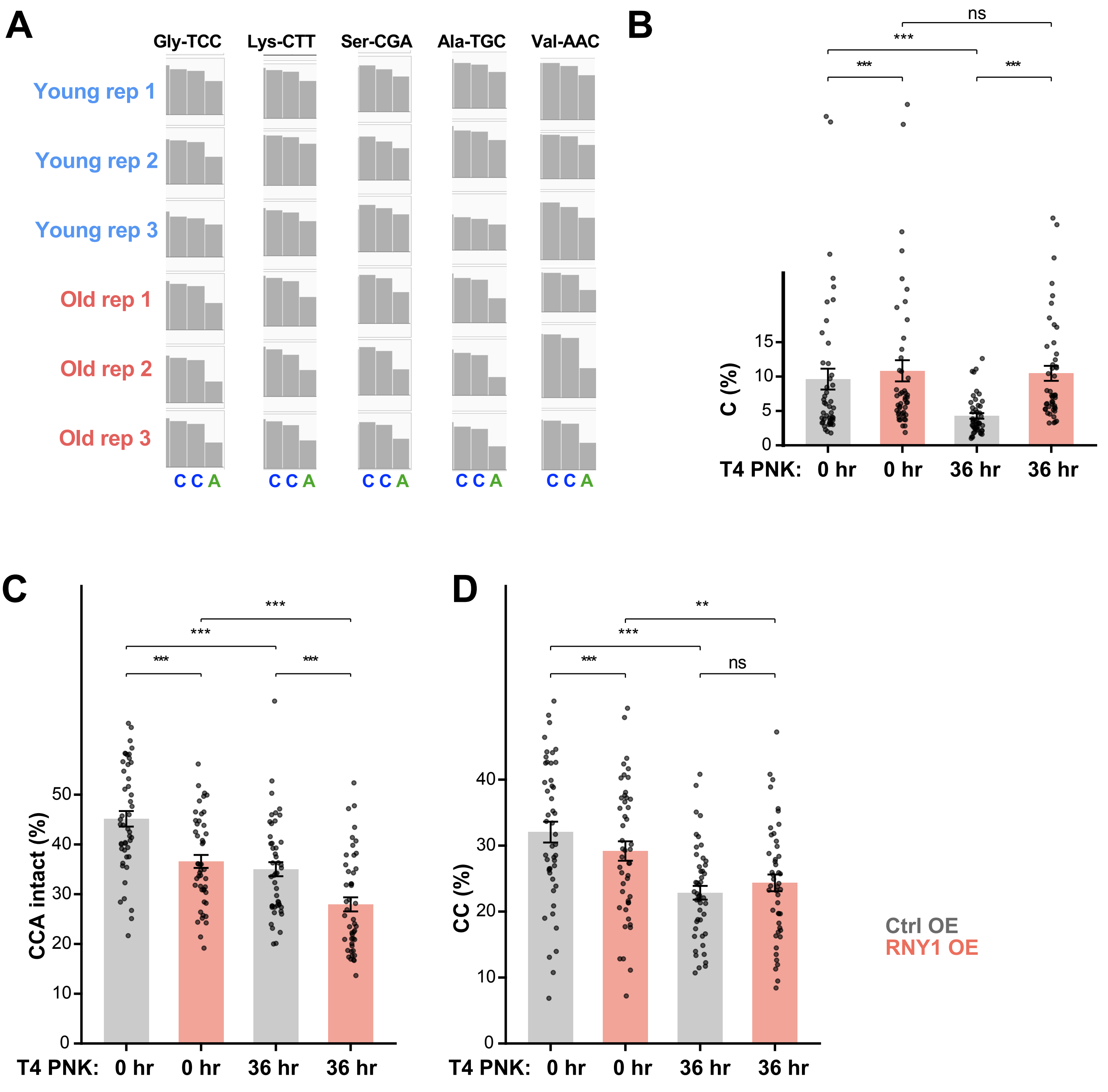
A
Gly-TCC
Lys-CTT
Ser-CGA
Ala-TGC
Val-AAC
Young rep 1
Young rep 2
Young rep 3
Old rep 1
Old rep 2
Old rep 3
C C A
B
ns
***
***
***
C (%)
15
10
5
0
T4 PNK: 0 hr 0 hr 36 hr 36 hr
C
CCA intact (%)
50
40
30
20
10
0
***
***
***
***
T4 PNK: 0 hr 0 hr 36 hr 36 hr
D
CC (%)
40
30
20
10
0
**
***
***
ns
T4 PNK: 0 hr 0 hr 36 hr 36 hr
Ctrl OE
RNY1 OE

### Figure S4

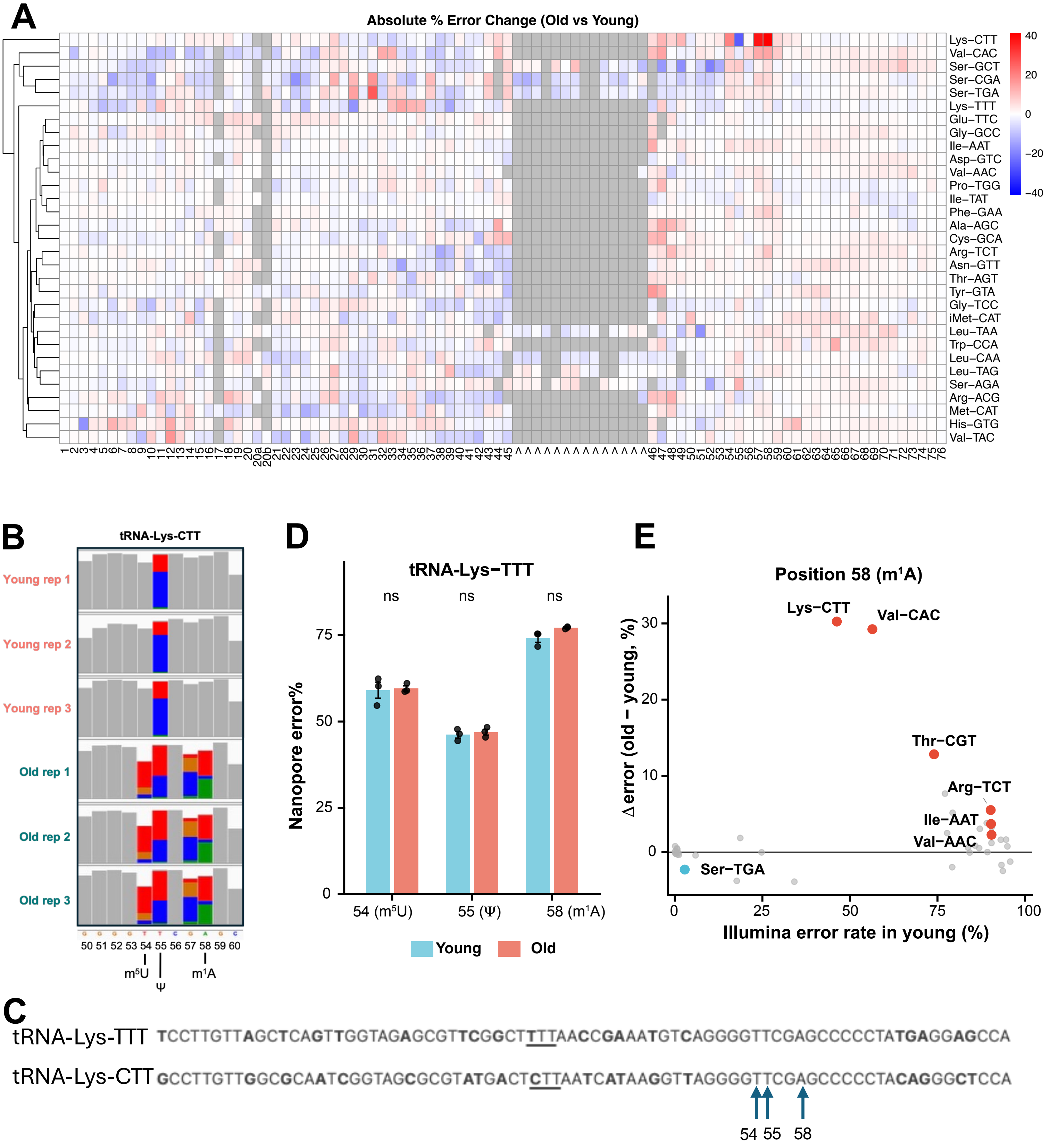
A
Absolute % Error Change (Old vs Young)
Lys–CTT
Val–CAC
Ser–GCT
Ser–CGA
Ser–TGA
Lys–TTT
Glu–TTC
Gly–GCC
Ile–AAT
Asp–GTC
Val–AAC
Pro–TGG
Ile–TAT
Phe–GAA
Ala–AGC
Cys–GCA
Arg–TCT
Asn–GTT
Thr–AGT
Tyr–GTA
Gly–TCC
iMet–CAT
Leu–TAA
Trp–CCA
Leu–CAA
Leu–TAG
Ser–AGA
Arg–ACG
Met–CAT
His–GTG
Val–TAC
40
20
0
−20
−40
B
tRNA-Lys-CTT
Young rep 1
Young rep 2
Young rep 3
Old rep 1
Old rep 2
Old rep 3
50 51 52 53 54 55 56 57 58 59 60
m5U
Ψ
m1A
D
tRNA−Lys−TTT
ns
ns
ns
Nanopore error%
0
25
50
75
54 (m5U)
55 (Ψ)
58 (m1A)
Young
Old
E
Position 58 (m1A)
Δ error (old − young, %)
Illumina error rate in young (%)
Lys−CTT
Val−CAC
Thr−CGT
Arg−TCT
Ile−AAT
Val−AAC
Ser−TGA
C
tRNA-Lys-TTT TCCTTGTTAGCTCAGTTGGTAGAGCGTTCGGCTTTTAACCGAAATGTCAGGGGTTCGAGCCCCCTATGAGGAGCCA
tRNA-Lys-CTT GCCTTGTTGGCGCAATCGGTAGCGCGTATGACTCTTAATCATAAGGTTAGGGGTTCGAGCCCCCTACAGGGCTCCA
54 55 58

### Figure S5

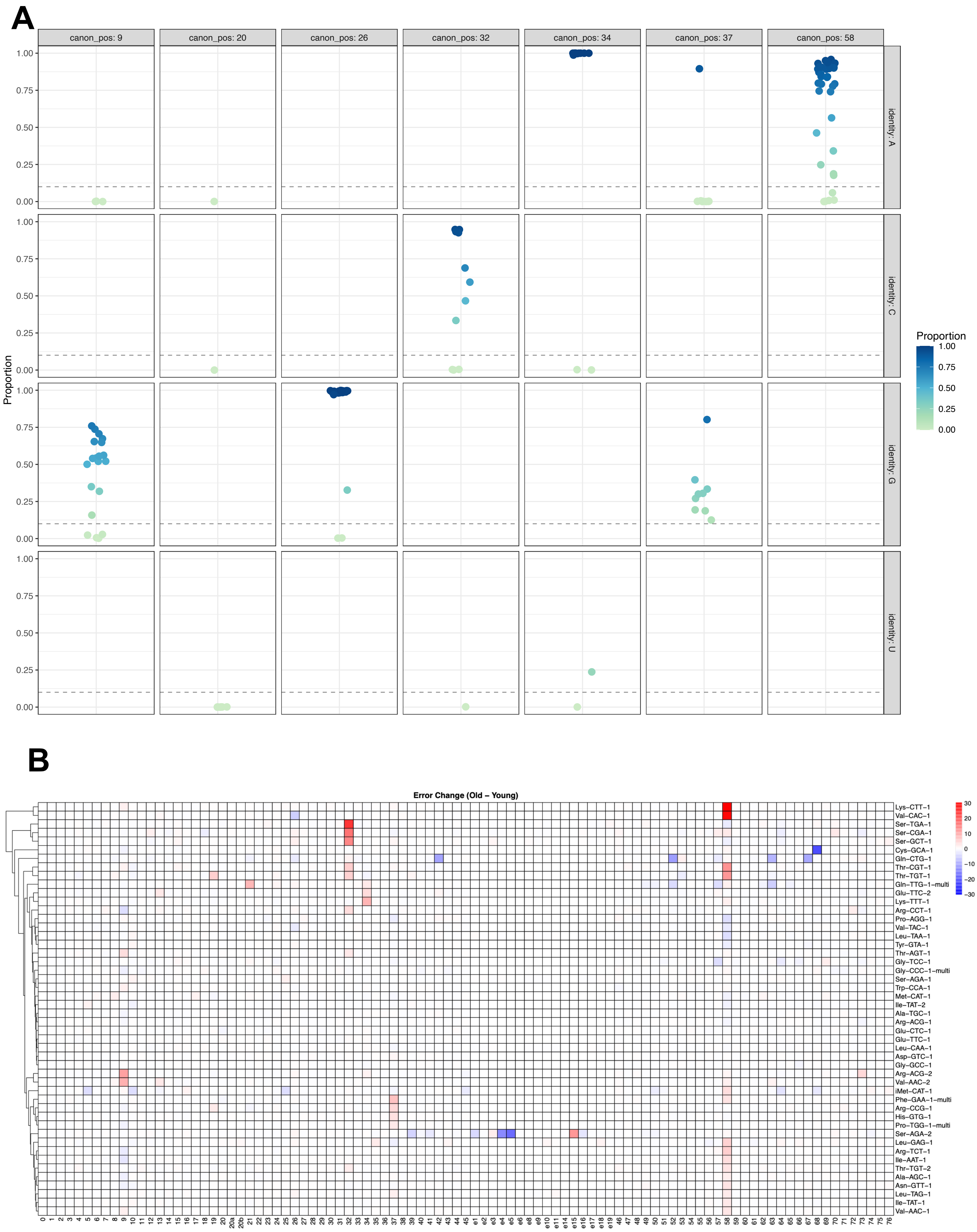
A
canon_pos: 9
canon_pos: 20
canon_pos: 26
canon_pos: 32
canon_pos: 34
canon_pos: 37
canon_pos: 58
identity: A
identity: C
identity: G
identity: U
Proportion
1.00
0.75
0.50
0.25
0.00
B
Error Change (Old – Young)
Lys–CTT–1
Val–CAC–1
Ser–TGA–1
Ser–CGA–1
Ser–GCT–1
Cys–GCA–1
Gln–CTG–1
Thr–CGT–1
Thr–TGT–1
Gln–TTG–1–multi
Glu–TTC–2
Lys–TTT–1
Arg–CCT–1
Pro–AGG–1
Val–TAC–1
Leu–TAA–1
Tyr–GTA–1
Thr–AGT–1
Gly–TCC–1
Gly–CCC–1–multi
Ser–AGA–1
Trp–CCA–1
Met–CAT–1
Ile–TAT–2
Ala–TGC–1
Arg–ACG–1
Glu–CTC–1
Glu–TTC–1
Leu–CAA–1
Asp–GTC–1
Gly–GCC–1
Arg–ACG–2
Val–AAC–2
iMet–CAT–1
Phe–GAA–1–multi
Arg–CCG–1
His–GTG–1
Pro–TGG–1–multi
Ser–AGA–2
Leu–GAG–1
Arg–TCT–1
Ile–AAT–1
Thr–TGT–2
Ala–AGC–1
Asn–GTT–1
Leu–TAG–1
Ile–TAT–1
Val–AAC–1
30
20
10
0
−10
−20
−30
